# Combined ADAMTS10 and ADAMTS17 inactivation exacerbates bone shortening and compromises extracellular matrix formation

**DOI:** 10.1101/2025.01.23.634616

**Authors:** Nandaraj Taye, Stylianos Z. Karoulias, Zerina Balic, Lauren W. Wang, Belinda B. Willard, Daniel Martin, Daniel Richard, Alexander S. Okamoto, Terence D. Capellini, Suneel S. Apte, Dirk Hubmacher

## Abstract

Weill-Marchesani syndrome (WMS) is characterized by severe short stature, short hands and feet (brachydactyly), joint contractures, tight skin, and heart valve, eye, and skin anomalies. Whereas recessive WMS is caused by mutations in *ADAMTS10*, *ADAMTS17*, or *LTBP2*, dominant WMS is caused by mutations in *FBN1* (encoding fibrillin-1). Since bone growth is driven by chondrocyte proliferation and hypertrophy in the growth plates, the genetics of WMS suggests that the affected ECM proteins act within the same pathway to regulate chondrocyte and growth plate function. Here, we investigated the role of the secreted ADAMTS proteases ADAMTS10 and ADAMTS17 in growth plate function and ECM formation. We generated *Adamts10*;*Adamts17* double knockout (DKO) mice, which showed significant postnatal lethality compared to single *Adamts10* or *Adamts17* KO mice. Importantly, we observed severe bone shortening DKO mice, which correlated with a narrower hypertrophic zone in their growth plates. ADAMTS17 substrates identified by N-terminomics and yeast two-hybrid screening identified the ECM proteins fibronectin and collagen VI (COL6). However, validation experiments did not reveal direct proteolysis of either fibronectin or COL6 by ADAMTS17. We then investigated ECM formation in primary ADAMTS10- and ADAMTS17-deficient skin fibroblasts and observed compromised fibronectin deposition concomitant with aberrant intracellular accumulation of fibrillin-1. These findings support a role for ADAMTS17 in ECM protein secretion and assembly. Collectively, our data suggest that ADAMTS10 and ADAMTS17 regulate bone growth by regulating chondrocyte hypertrophy or hypertrophic chondrocyte turnover. Mechanistically, ADAMTS17 appears to be a critical regulator of ECM protein secretion or pericellular matrix assembly, whereas ADAMTS10 likely modulates ECM formation at later stages, possibly regulating the spatio-temporal deposition of fibrillin isoforms.

## Introduction

Acromelic dysplasias are a group of genetic conditions resulting in severe short stature and shortening of distal limb elements^1,2^. Among them, Weill-Marchesani syndrome (WMS) has been characterized genetically resulting from mutations in distinct, but functionally related genes, encoding extracellular matrix (ECM) proteins^1,3^. Recessive pathogenic variants in secreted ADAMTS10 cause Weill-Marchesani syndrome 1 (WMS1)^4–6^. Distinct WMS sub-types can also be caused by dominant mutations in fibrillin-1 (*FBN1*, WMS2) or recessive mutations in *LTBP2* (WMS3) or *ADAMTS17* (WMS4) suggesting that these four genes may act together in pathways that regulate development and homeostasis of affected tissues^4,7–11^. WMS is characterized by short stature, lens dislocation, microspherophakia and other eye anomalies, progressive joint stiffness, and tight skin^3,12^. Lens dislocation and/or changes in the outflow track can block the drainage of aqueous humor from the anterior chamber of the eye and cause glaucoma in WMS. WMS1 also has cardiovascular manifestations such as patent ductus arteriosus, pulmonary valve dysplasia, which can lead to pulmonary stenosis, and in rare cases aortic aneurysms^6,8,12^. In dogs, mutations in *ADAMTS10* and *ADAMTS17* were associated with primary open angle glaucoma^13–17^. Homozygosity of a glaucoma-causing canine *ADAMTS17* variants was also associated with short stature in several dog breeds^18^. WMS-causing ADAMTS10 and ADAMTS17 mutations are distributed over the entire molecule and result in loss-of-function or haploinsufficiency due to impaired secretion^1,3,10,11^. The fact that mutations in *ADAMTS10* and *ADAMTS17* each cause WMS suggests that these genes cooperate, have superimposed mechanisms, or act in the same pathways that regulate the development or homeostasis of affected tissues, including the growth plate, which drives bone growth, the skin, and the eye^19,20^. ADAMTS10 and ADAMTS17 each bind to fibrillin-1, which may provide a scaffold for their tissue-specific deposition or functional regulation^21,22^. In addition, ADAMTS10 was shown to promote the assembly of fibrillin-1 in cell culture^21^.

The 19 ADAMTS proteases are involved in diverse biological processes including tissue morphogenesis and homeostasis^23,24^. Mutations in several ADAMTS proteases cause birth defects and inherited connective tissue disorders in humans and other species^25^. In addition, ADAMTS proteases contribute to the progression of acquired disease, such as the aggrecanase ADAMTS5 in osteoarthritis^26^. These diverse roles are attributed to the actions of ADAMTS proteases on distinct substrates. Recognized ECM substrates for ADAMTS proteases include proteoglycans such as aggrecan and versican, the ECM scaffolding proteins fibrillin-1, fibrillin-2, and fibronectin, and several collagens^23^. Most ADAMTS proteases are secreted as inactive zymogens whose activation requires proteolytic removal of their prodomains by furin or other proprotein convertases^27–30^. ADAMTS10, however, lacks a canonical furin recognition site and is poorly processed by furin. Consistent with poor furin-processing, ADAMTS10 appears to be an inefficient protease, although, once activated by mutagenesis to restore a canonical furin-processing site, ADAMTS10 could cleave fibrillin-1 and fibrillin-2 in vitro^21,31^. However, alternative furin-independent mechanisms of ADAMTS10 activation in vitro or in vivo remain elusive. ADAMTS17 is secreted as an active protease, but fragments itself extensively and efficiently, including within the catalytic domain, prior to its release from the surface of HEK293 cells^22^. These findings suggested that ADAMTS17 may act as a protease in the secretory pathway or at the cell surface. ADAMTS17 substrates other than itself have not been identified. We showed previously that ADAMTS17 interacted with fibrillin-1 and fibrillin-2 but did not cleave either^22,32^.

*Adamts10* and *Adamts17* inactivation in mice was previously reported with differential phenotypes. While *Adamts10* knockout (KO) mice did not result in short stature, *Adamts17* KO mice had shorter bones and growth plate abnormalities^31,33^. Interestingly, the knock-in of a WMS mutation into the mouse *Adamts10* locus resulted in short stature and growth plate abnormalities^34^. In addition, knock-in of an *ADAMTS10* mutation that causes glaucoma in dogs into the mouse *Adamts10* locus also resulted in short stature^35^. Together, these findings support a role for ADAMTS10 and ADAMTS17 in regulating bone growth and thus height. If and how ADAMTS10 and ADAMTS17 cooperate in regulating bone growth and in the formation and maintenance of other tissues affected in WMS is not known. Here, we investigated the genetic interactions of ADAMTS10 and ADAMTS17 by analyzing the bone and skin phenotypes of *Adamts10*;*Adamts17* double KO (DKO) mice. For insights on molecular mechanisms, we evaluated potential ADAMTS17 substrates and binding partners identified by N-terminomics and yeast two-hybrid screening, respectively. The findings of our studies, taken together with prior work strongly suggest a cooperative role for these proteases in skeletal growth and provide a putative molecular basis for their ECM-regulatory activities.

## Results

### Combined *Adamts10* and *Adamts17* inactivation resulted in early postnatal mortality and reduced body size

To generate *Adamts10*;*Adamts17* DKO mice, we combined a previously published *Adamts10* KO allele with a novel *Adamts17* KO allele. The *Adamts10* KO allele was generated by replacing 41 bp of *Adamts10* exon 5 with an IRES-*lacZ*-*neo* cassette^31^. The *Adamts17* KO allele was generated using CRISPR/Cas9-induced non-homologous end-joining with guide RNAs targeting *Adamts17* exon 3 (Fig. 1A). The resulting dinucleotide insertion (AT) in exon 3 of *Adamts17* (*Adamts17* 670_671insAT, NM_001033877.4) caused a reading frame shift (p.I190fsX12), which resulted in a premature termination codon presumed to trigger nonsense-mediated mRNA decay (Fig. 1A). The AT insertion in *Adamts17* was verified by Sanger sequencing of a PCR product amplified from template DNA isolated from toe tissue with *Adamts17*-specific primers flanking exon 3 (Fig. 1B). Mice homozygous for the *Adamts17* AT insertion are referred to as *Adamts17* KO. Loss of ADAMTS17 protein was validated by immunostaining tissue and primary skin fibroblasts isolated from wild type (WT) and *Adamts17* KO mice using a monoclonal ADAMTS17 antibody. In sections through WT skin and the tibial growth plate, ADAMTS17 immunoreactivity was apparent around hair follicles and in hypertrophic chondrocytes, respectively. This signal was absent or strongly reduced in comparable sections from *Adamts17* KO or *Adamts10*;*Adamts17* DKO mice (Fig. 1C). In primary WT skin fibroblasts, the ADAMTS17 signal was most intense in perinuclear regions, where the endoplasmic reticulum and Golgi are located, and in patches between cells (Fig. 1C, right). Similar to tissue sections, the ADAMTS17 signal was absent in DKO fibroblasts. Thus, genetic inactivation of ADAMTS17 eliminated ADAMTS17 protein in relevant tissues and primary cells and at the same time validated the specificity of the monoclonal ADAMTS17 antibody.

**Figure 1.**
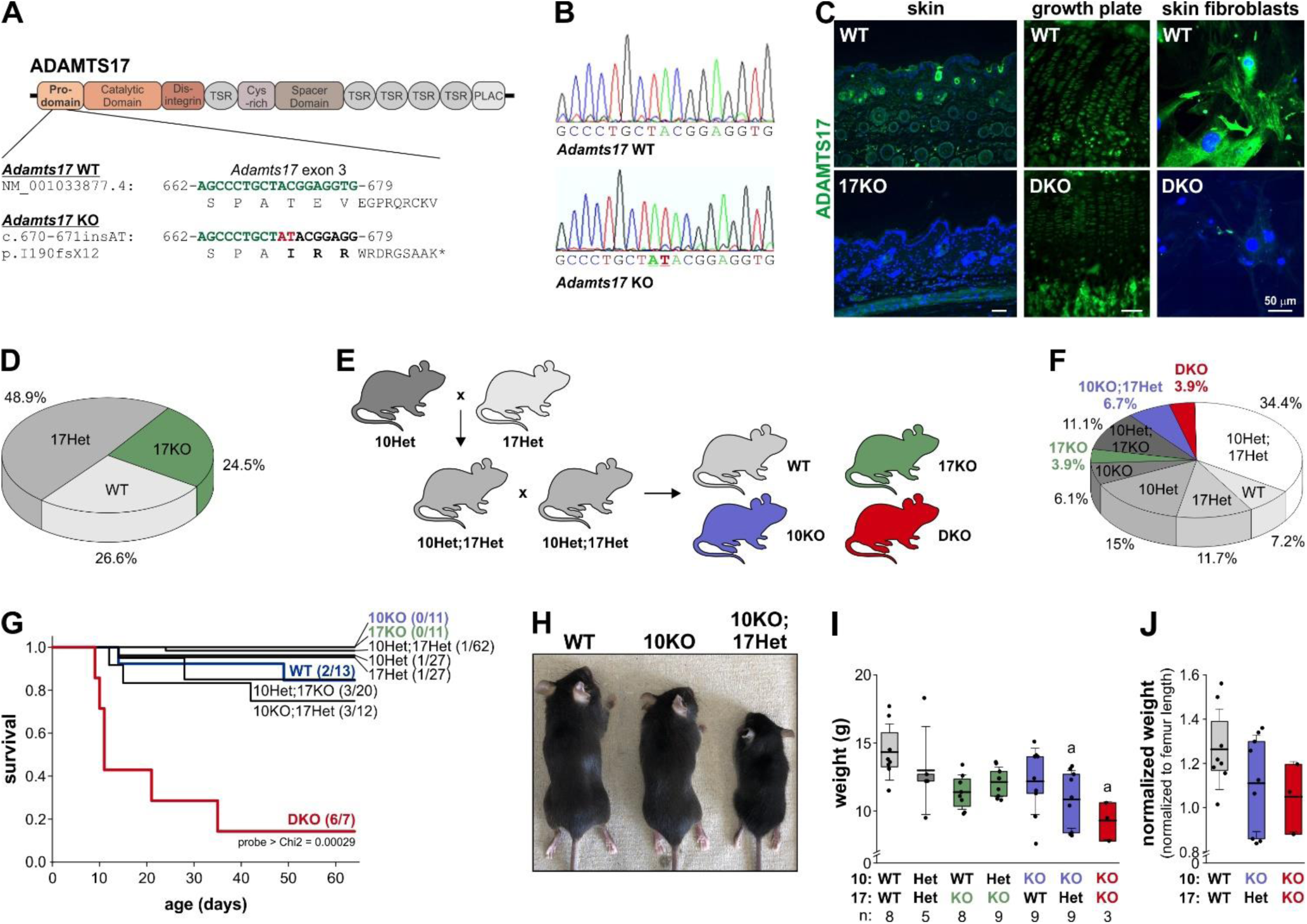
Generation of Adamts17 KO and Adamts10;Adamts17 DKO mice. **A)** Domain organization of ADAMTS17 shows location and targeting of exon 3 by CRISPR/Cas9 gRNA to induce non-homologous end joining. The nucleotide and amino acid sequence of the ADAMTS17 WT allele (green) and after AT insertion (red) are indicated. The dinucleotide insertion induced a frameshift, which resulted in a premature stop codon after 12 amino acids. **B)** Sanger sequencing traces of a PCR product generated with primers flanking exon 3 showing the AT insertion (underlined) in the Adamts17 KO. **C)** Micrographs of ADAMTS17 immunostaining of sections through WT and Adamts17 KO skin (left), DKO growth plates (middle), and of primary DKO mouse skin fibroblasts (right). The signal in the dermis around hair follicles, in growth plate chondrocytes, and in fibroblasts and their ECM originating from the monoclonal ADAMTS17 antibody was strongly reduced in KO and DKO tissues and cells, indicating lack of ADAMTS17 protein in Adamts17 KO mice. **D)** Pie chart showing Mendelian distribution of genotypes recovered from Adamts17 Het intercrosses at the time of genotyping (P7-P10) (n=94 mice). **E)** Breeding scheme to generate WT, Adamts10 KO (10KO), Adamts17 KO (17KO), and DKO mice. **F)** Pie chart showing distribution of genotypes recovered from Adamts10 Het;Adamts17 Het intercrosses at P7-P10 (n=180 mice). Statistical analysis was performed using Chi square calculation. **G)** Kaplan-Meier survival analysis of DKO mice. The numbers of observed dead/total mice for the individual genotypes are indicated in brackets. Statistical significance was determined using a log-rank test. **H)** Whole mount images of WT, 10KO, 10KO;17Het mice at 4 weeks of age shows progressive reduction in body size. **I)** Bar graphs showing body weights of 4-week-old mice of the indicated genotypes. The number of mice is indicated below the genotypes. **J)** Bar graphs showing body weight normalized to average femur length for the genotypes that were significantly different in I. In I, J floating bars indicate the 25^th^ – 75^th^ percentile range, lines the mean value, and whiskers the standard deviation. Statistical differences in I, J were determined using a one-way ANOVA with post-hoc Tukey test. a, p<0.05 compared to WT.

*Adamts17* KO mice were born at the expected Mendelian ratio and were viable (Fig. 1D). To generate DKO mice, we first crossbred *Adamts10* Het and *Adamts17* Het mice to generate *Adamts10;Adamts17* double-heterozygous mice (Fig. 1E). *Adamts10* Het;*Adamts17* Het mice were then intercrossed to generate offspring with all allelic combinations, among which WT, *Adamts10* KO; *Adamts17* KO, and *Adamts10*;*Adamts17* DKO mice were the focus of subsequent analyses. In principle, this breeding scheme allows comparison of the phenotypes from littermates. However, due to the low predicted percentages for WT and DKO mice (6.25%) per litter in these crosses, we also intercrossed *Adamts10* Het or *Adamts17* Het mice to generate the respective WT and individual KOs as age- and sex-matched controls for DKO mice. We first analyzed the *Adamts10* and *Adamts17* genotype distribution of 180 mice at postnatal day (P) 7-P10 and determined statistically significant deviations from the expected Mendelian ratios using Chi^2^ calculation (Fig. 1F). *Adamts17* KO (3.9% vs. 6.25 expected), *Adamts10* KO;*Adamts17* Het (6.7% vs 12.5% expected), and DKO (3.9% vs. 6.25 expected) mice were present in significantly lower numbers than expected and the percentage of *Adamts10*;*Adamts17* double heterozygous mice was significantly higher (34.4% vs. 25% expected). This resulted in a Chi^2^ value of 18.09 (8 degrees of freedom) and a p-value of <0.05, suggesting reduced viability or embryonic lethality due to reduced *Adamts17* gene dosage or the combined absence of *Adamts10* and *Adamts17*. Since we observed early postnatal lethality of *Adamts10*;*Adamts17* DKO mice, we quantified postnatal survival with Kaplan-Meier survival analysis, where we observed significant postnatal mortality of DKO mice with 50% survival at 23 d (+/-7.1 d) after birth (probe > Chi^2^ = 0.00029, log-rank test) (Fig. 1G). The cause of death is unknown. In addition to reduced survival of DKO mice, we noted reduced body size, which correlated with several genotypes, most notably *Adamts17* KO mice, *Adamts10* KO;*Adamts17* Het mice and DKO mice (Fig. 1H). These size differences were also apparent from body weight measurements at 4 weeks of age where the weights of *Adamts10* KO;*Adamts17* Het and DKO mice were significantly reduced compared to WT (Fig. 1I). However, after normalization of the body weight to the average femur lengths, these differences became non-significant, suggesting proportionate short stature (Fig. 1J).

### Combined *Adamts10*;*Adamts17* depletion exacerbated bone shortening

To determine if *Adamts10* and *Adamts17* gene dosage affected bone length, we quantified the lengths of forelimb and hind limb bones from X-ray images taken at 4 weeks of age (Fig. 2A-L). Overall, *Adamts10*;*Adamts17* DKO mice had the shortest bones across all genotypes when compared to WT or individual KOs. In addition, *Adamts17* KO bones were significantly shorter than WT, except for the humerus, the 1^st^ metacarpal, and the 1^st^ metatarsal. Notably, deletion of one *Adamts17* allele in *Adamts10* KO mice exacerbated bone shortening, but not vice versa, except in the fibula, the 1^st^ metacarpal, and the 1^st^ metatarsal. The lengths of the 1^st^ metacarpal and 1^st^ metatarsal were significantly shorter in DKO mice compared to WT, *Adamts10* KO, and *Adamts17* KO but the same bones were not shorter in the individual KOs compared to WT (Fig. 2E, L). The lengths of *Adamts10;Adamts17* double heterozygous bones were not significantly shorter compared to WT bones. Since we previously observed increased bone width in another acromelic dysplasia model (geleophysic dysplasia) due to *Adamtsl2* deficiency, we measured the width of humeri and femora^36^. The mid-shaft width in DKO mice was significantly greater than WT, *Adamts10* KO, or *Adamts17* KO (humerus), or compared to WT (femur) (Fig. 2M, N). In addition, the widths of the *Adamts10* KO;*Adamts17* Het humerus and femur were increased compared to WT, but not the individual KOs. Collectively, combined inactivation of ADAMTS10 and ADAMST17 exacerbated bone shortening compared to the individual KOs with *Adamts17* having an apparently stronger gene dosage effect compared to *Adamts10*.

**Figure 2.**
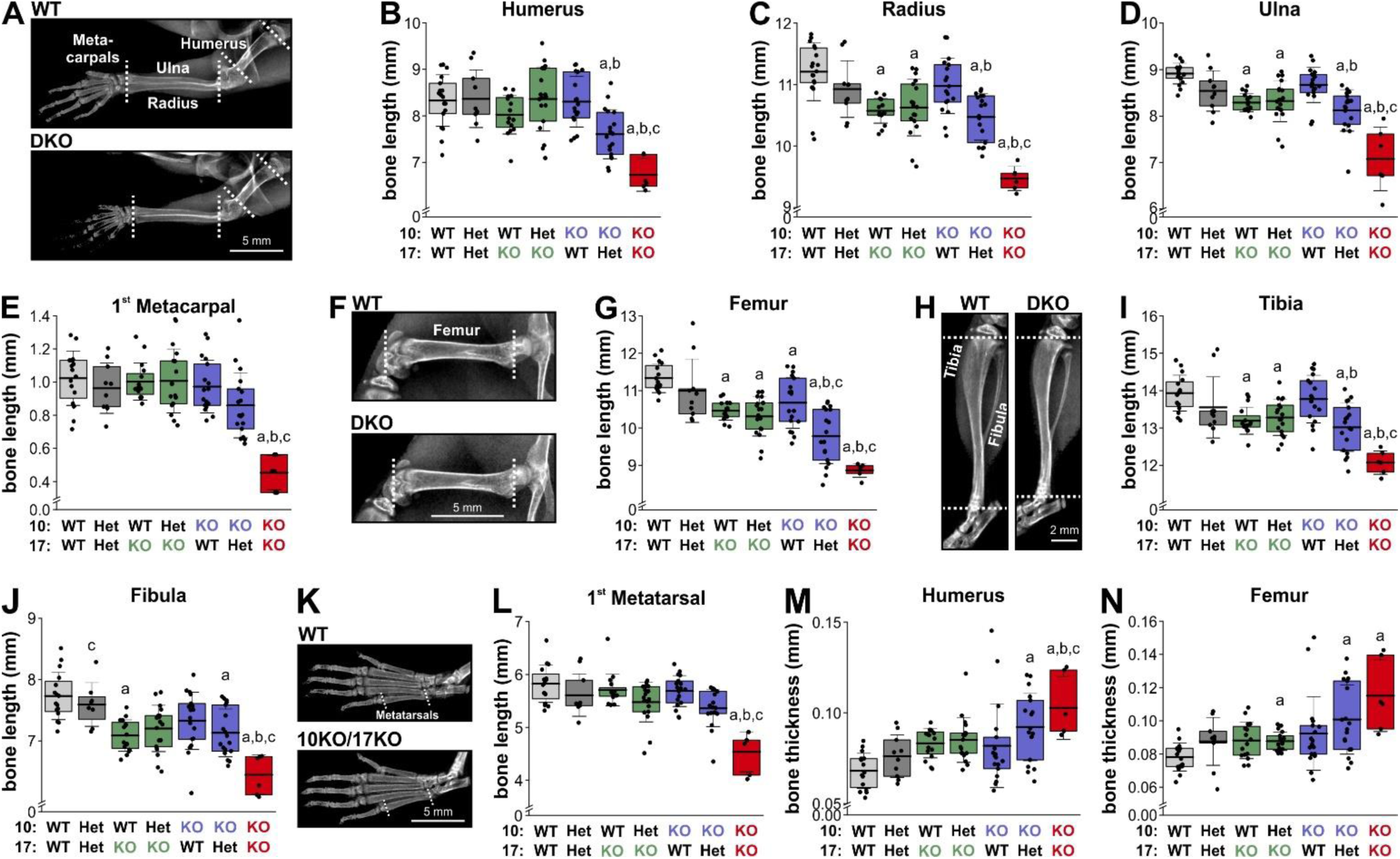
Exacerbated bone shortening in Adamts10;Adamts17 DKO mice. **A)** X-ray images showing WT and DKO forelimbs. **B-E)** Bar graphs showing lengths of humerus (B), radius (C), ulna (D), and 1^st^ metacarpal (E) for all genotypes. **F)** X-ray images showing WT and DKO femur. **G)** Bar graphs showing femoral length for all genotypes. **H)** X-ray images showing WT and DKO tibia and fibula. **I, J)** Bar graphs showing lengths of tibia (I) and fibula (J) for all genotypes. **K)** X-ray images showing WT and DKO hind paw bones. **L)** Bar graphs showing length of 1^st^ metatarsal for all genotypes. **M, N)** Bar graphs showing thickness of humerus (M) and femur (N) at mid-shaft. For number of mice see Fig. 1I. All mice were 4 weeks old at the time of X-ray imaging. Bones from both limbs were measured and measurement from male and female mice were combined. In B, C, D, E, G, I, J, L, M, N floating bars indicate the 25^th^ – 75^th^ percentile range, lines the mean value and whiskers the standard deviation. Statistical differences in B, C, D, E, G, I, J, L, M, N were determined using a one-way ANOVA with post-hoc Tukey test. a, p<0.05 compared to WT; b, p<0.05 compared to Adamts10 KO; c, p<0.05 compared to Adamts17 KO.

### Combined *Adamts10* and *Adamts17* inactivation compromised growth plate chondrocyte function

Since bone growth is largely driven by growth plate activity, i.e. the proliferation and hypertrophic expansion of growth plate chondrocytes, we quantified growth plate dimensions and primary chondrocyte behavior in ADAMTS10 and ADAMTS17 deficient tissue and chondrocytes^37,38^. Compared to WT growth plates, DKO growth plates showed a narrower hypertrophic zone, while the proliferative zone was unchanged (Fig. 3A, B). In addition, the columnar organization of proliferating chondrocytes appeared to be irregular with more spacing between chondrocyte columns (Fig. 3C). By immunostaining, we localized ADAMTS17 in the pericellular matrix of hypertrophic chondrocytes in wild type growth plates, close to the cartilage-bone interface (Fig. 3D). These data suggested that ADAMTS17 could play a role in regulating or maintaining chondrocyte hypertrophy. Therefore, we next used high cell density pellet cultures of mouse embryo CH3/10T1/2 cells to model chondrocyte-like differentiation in vitro and to determine the temporal dynamics of *Adamts10* and *Adamts17* mRNA expression during chondrocyte maturation in vitro by quantitative real-time (qRT)-PCR (Fig. 3E). *Adamts10* mRNA levels significantly decreased between 0 and 2 days after induction of differentiation and remained low thereafter (Fig. 3F). In contrast, *Adamts17* mRNA levels started to moderately increase until 7 days after induction of differentiation followed by a strong and statistically significant increase at 10 days. This suggested downregulation of *Adamts10* and induction of *Adamts17* gene expression during in vitro chondrogenesis of CH3/10T1/2 pellet cultures. The strong reduction of *Col2a1* mRNA levels between 0 and 2 days after induction of differentiation was consistent with CH3/10T1/2 pellet cultures undergoing differentiation.

**Figure 3.**
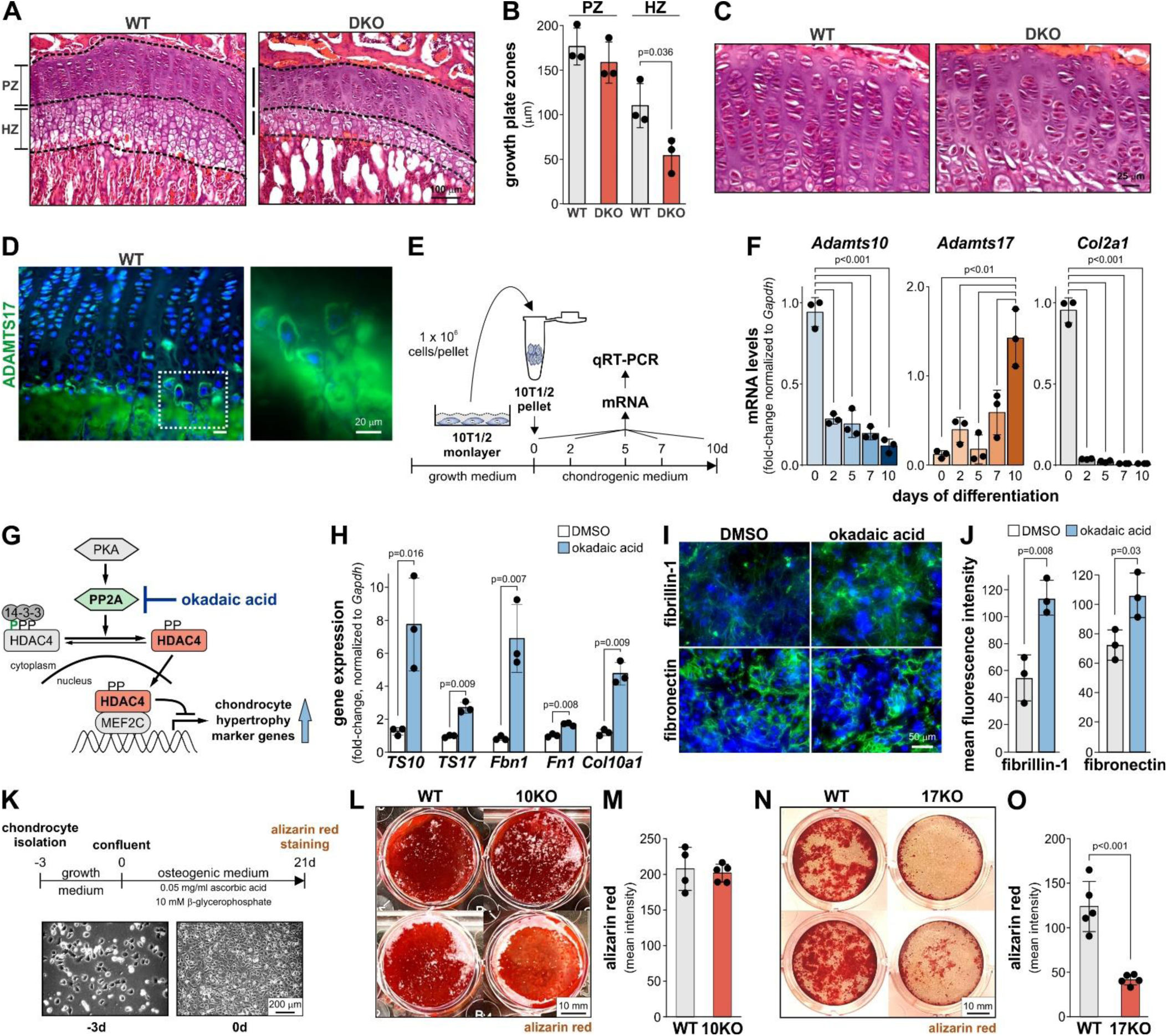
ADAMTS10 and ADAMTS17 regulate growth plate function and chondrocyte hypertrophy. **A)** Images of sections through growth plates of 4 week-old WT and Adamts10;Adamts17 DKO mice. Proliferative (PZ) and hypertrophic (HZ) zones are outlined with dashed lines. **B)** Bar graphs showing widths of PZ and HZ from WT and DKO growth plates. Data points represent the average of multiple measurements across the growth plate zone from n=3 mice. **C)** Higher magnification of growth plate from images in A showing disorganized proliferative zone in DKO growth plates. **D)** Micrograph of ADAMTS17 immunostaining of hypertrophic chondrocytes at the cartilage-bone interface. The boxed area is magnified in the right-hand panel. **E)** Schematic representation of experimental design for pellet culture to induce chondrocyte-like differentiation of C3H/10T1/2 cells. **F)** Bar graphs showing relative Adamts10, Adamts17, and Col2a1 mRNAs levels during differentiation of C3H/10T1/2 cell pellets normalized to Gapdh (n=3 replicates). **G)** Schematic representation of the mechanism of action of okadaic acid in de-repressing chondrocyte hypertrophy genes. **H)** Bar graphs showing relative changes of Adamts10 (TS10), Adamts17 (TS17), Fbn1, Fn1, and Col10a1 mRNA levels 24 h after treatment of primary chondrocytes with 50 nM okadaic acid or DMSO (n=3 replicates). **I)** Micrographs of immunostaining of fibrillin-1 (FBN1) and fibronectin (FN) deposition in the ECM of primary chondrocytes 3 days after treatment with okadaic acid or DMSO only. **J)** Quantification of mean fluorescence intensity from I (n=3 fields-of-view). **K)** Schematic representation of osteogenic differentiation of P5 primary rib chondrocytes isolated from Adamts10 KO or Adamts17 KO mice. The bottom panels show brightfield micrographs of freshly isolated primary chondrocytes (-3 d, left) and confluent chondrocytes (0 d, right). **L)** Micrographs of two individual wells/genotype of primary WT or Adamts10 KO (10KO) chondrocytes stained with alizarin red after 21 d of culture in osteogenic medium. **M)** Bar graph showing quantification of mean signal intensity of alizarin red deposits (isolates from n=4-5 biological replicates/genotype). **N)** Micrographs of two individual wells/genotype of primary WT or Adamts17 KO chondrocytes stained with alizarin red after 21 d of culture in osteogenic medium. **O)** Bar graph showing quantification of mean signal intensity of alizarin red deposits (isolates from n=5 biological replicates/genotype). In B, F, H, J, M, O, bars indicate mean values and whiskers the standard deviation. Statistical significance in B, H, J, M, O was calculated with a 2-sided Student t-test and in F with one-way ANOVA followed by post-hoc Tukey test.

To further probe the regulation of *Adamts10* and *Adamts17* expression during chondrocyte hypertrophy, we treated primary WT rib chondrocytes with okadaic acid, a protein phosphatase 2A inhibitor, which results in de-repression of chondrocyte hypertrophy genes (Fig. 3G)^39,40^. qRT-PCR showed significant upregulation of *Adamts10* and *Adamts17* mRNA levels 24 h after treatment of primary WT chondrocytes with okadaic acid compared to DMSO-treated control chondrocytes (Fig. 3H). Increased *Col10a1* mRNA levels, a marker for hypertrophic chondrocytes, served as a positive control for okadaic acid-mediated de-repression of chondrocyte hypertrophy genes. In addition to *Adamts10* and *Adamts17*, mRNAs for genes encoding the ECM proteins fibrillin-1 (*Fbn1*) and fibronectin (*Fn1*) were also induced. Fibrillin-1 binds to ADAMTS10 and ADAMTS17 and mutations in *FBN1* cause WMS2^9,21,22^. Fibronectin forms the ECM scaffold required for fibrillin-1 deposition in the ECM of mesenchymal cells^41^. Using immunostaining, we confirmed increased fibrillin-1 and fibronectin ECM deposition in primary WT chondrocytes following okadaic acid treatment (Fig. 3I, J).

Finally, we investigated the implications of ADAMTS10 and ADAMTS17-deficiency on primary rib chondrocyte hypertrophy and the capacity to deposit calcium as hydroxyapatite in their ECM, which is a characteristic of terminal hypertrophic chondrocytes (Fig. 3K). Chondrocytes isolated from *Adamts10* KO ribs showed no difference in alizarin red-positive calcium mineral deposition after 21 d under differentiation and mineralization conditions (Fig. 3L, M). In contrast, calcium mineral deposition by ADAMTS17-deficient primary chondrocytes was significantly reduced, suggesting differential roles or differential compensation for ADAMTS10 and ADAMTS17 in chondrocyte differentiation (Fig. 3N, O).

To gain further insights into the epigenetic and transcriptomic regulation of *ADAMTS10* and *ADAMTS17* during human embryonic development, we data-mined recent assays for transposase-accessible chromatin with sequencing (ATAC-seq) and RNA-transcriptomic data sets, generated from micro- dissected cartilaginous human fetal appendicular skeletal elements (Fig. 4A)^42^. We first identified open chromatin regions by ATAC-seq within +/- 100 kb of *ADAMTS10* and *ADAMTS17* (Fig. 4B, C); each interval containing regulatory elements, such as enhancers, promoters, and repressor sequences, likely drives expression of the nearby gene. For *ADAMTS10*, out of ten total elements within 100 kbp, we identified three cartilage open chromatin regions that were shared by most or all autopod elements (phalanges, metatarsals and metacarpals), but were absent in stylopod or zeugopod elements, i.e. proximal and distal ends of all major long bones (Fig. 4B, Table 1). One regulatory region was located 100 kb distal to the ADAMTS10 transcription start site (TSS), one overlapping with the final exon, and the other <1 kb proximal to the TSS. We identified two additional cartilage open chromatin regions present in all skeletal elements, located within the *ADAMTS10* gene body. For *ADAMTS17*, we observed a larger number of regulatory elements compared to *ADAMTS10,* consistent with the larger size of *ADAMTS17*. At this locus, 23 cartilage open chromatin regions were identified within or in close vicinity to the *ADAMTS17* gene body (Fig. 4C, Table 2). While three *ADAMTS17* open chromatin regions were identified as accessible in all autopod elements and eight in most, two were specific to individual skeletal elements of the hind limb autopod (metatarsal V). Moreover, in the stylopod and zeugopod elements, two open chromatin regions were shared by the proximal elements but showed mixed accessibility in the corresponding distal ones.

**Figure 4:**
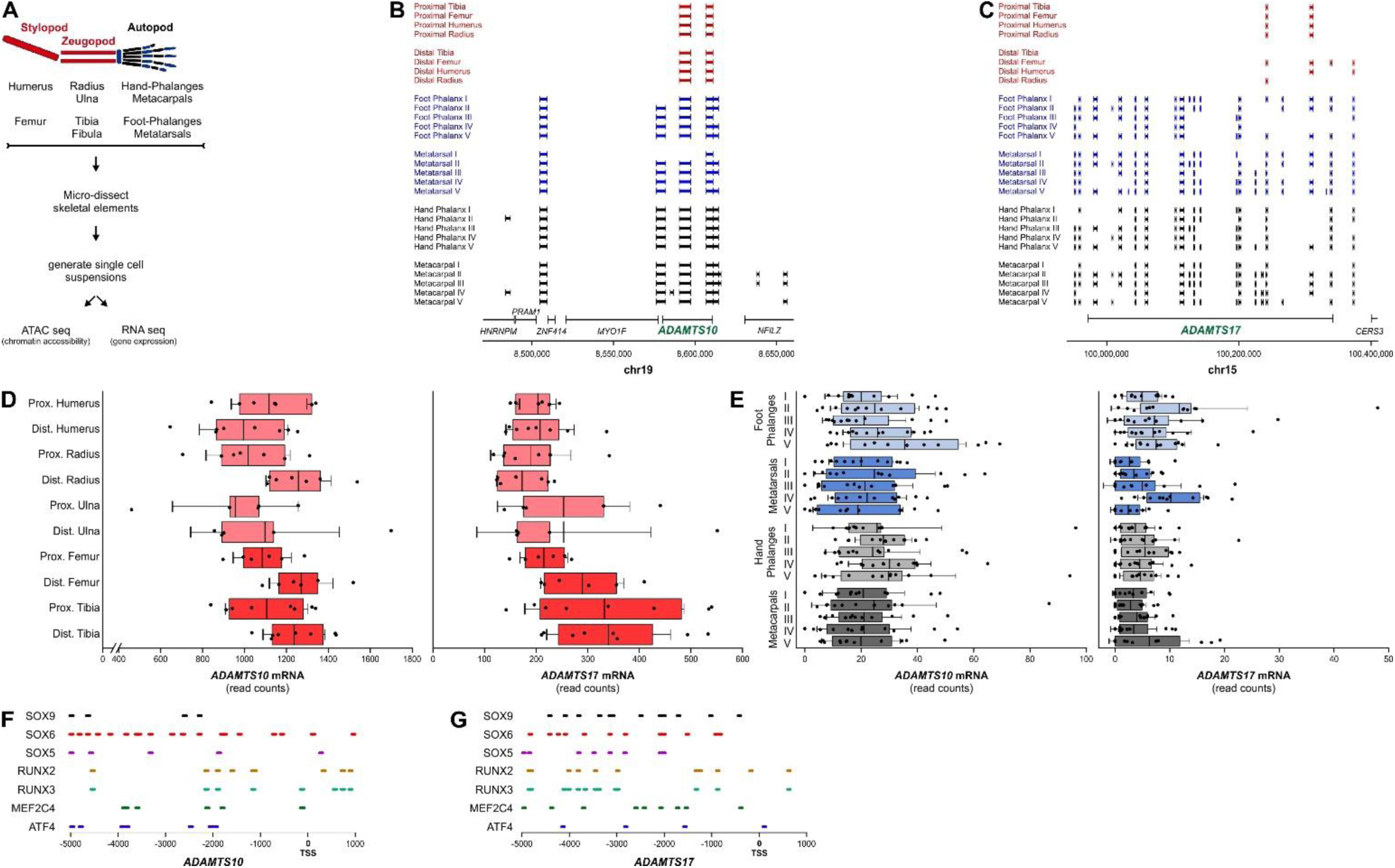
Chromatin accessibility and putative regulation of ADAMTS10 and ADAMTS17 in human cartilage and bone development. **A)** Limb skeletal elements and experimental design to generate ATAC-seq and transcriptomics data from skeletal elements of human products from conception (E54&E67). **B, C)** Mapping of open chromatin regions identified by ATAC-seq 100 kb up- or downstream of ADAMTS10 (B) and ADAMTS17 (C). The positions of ADAMTS10 on human chromosome 19 and ADAMTS17 on human chromosome 15 are indicated. **D, E)** Normalized read counts indicating ADAMTS10 and ADAMTS17 mRNA abundance in skeletal elements from stylopods and zeugopods (D) and autopods (E). **F, G)** Location of transcription factor binding sites 5 kb upstream of the ADAMTS10 (F) or ADAMTS17 (G) transcriptional start site (TSS).

**Table 1:**
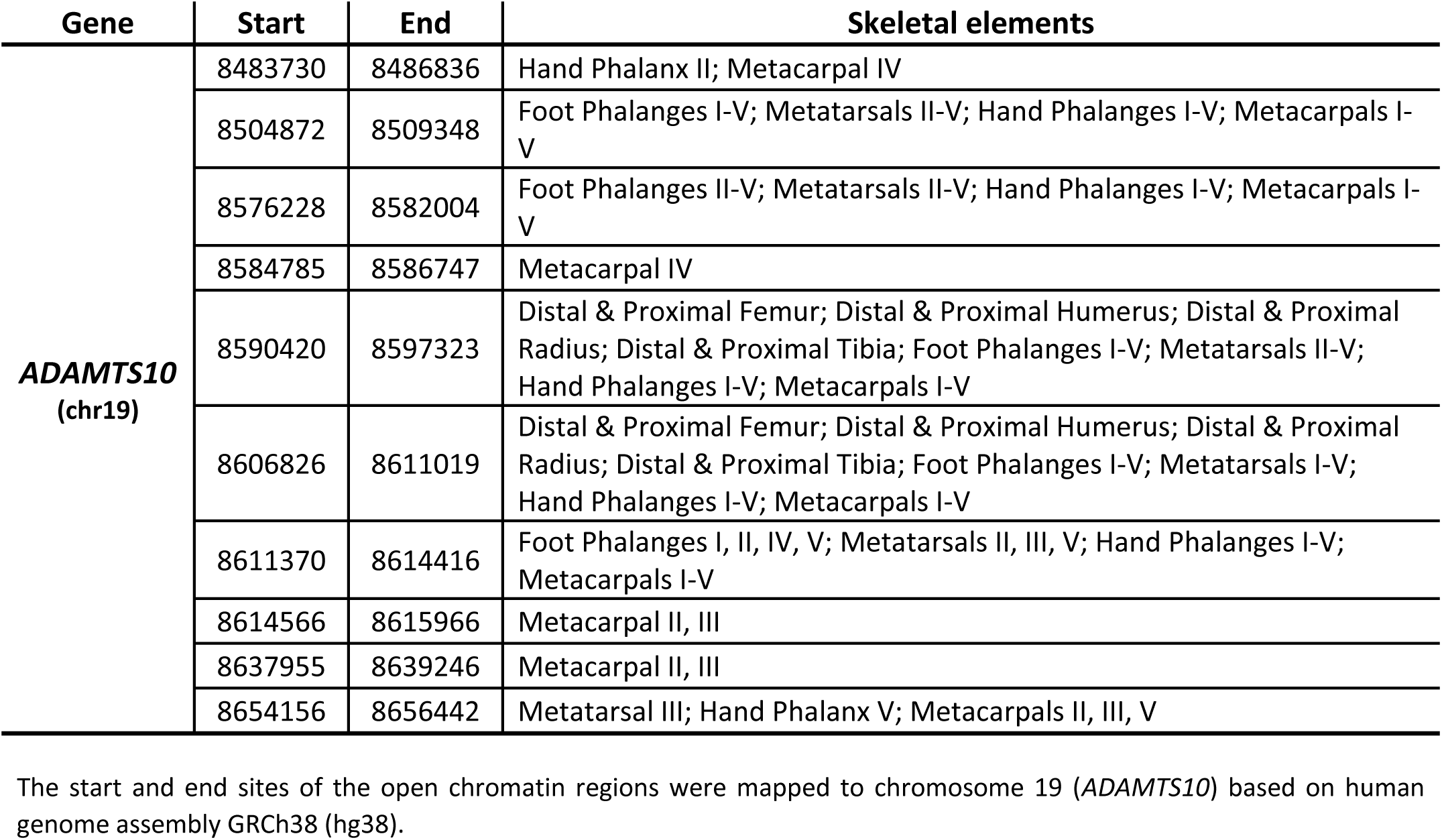
Accessible genomic regions in *ADAMTS10* in cartilage as determined by ATAC-seq.

**Table 2:** Accessible genomic regions in *ADAMTS17* in cartilage as determined by ATAC-seq.

| Gene | Start | End | Skeletal elements |
| --- | --- | --- | --- |
| ***ADAMTS17***<br>(chr15) | 99950945 | 99953010 | Foot Phalanges II-V; Metatarsals I-V; Hand Phalanges III-V; Metacarpals II-V |
|  | 99957622 | 99960241 | Foot Phalanges I-III, V; Metatarsals I-V; Hand Phalanges I, III, IV, V; Metacarpals I-III, V |
|  | 99980194 | 99985410 | Foot Phalanges I-III, V; Metatarsals I, II, V; Hand Phalanges III, V; Metacarpals II, III, V |
|  | 100007670 | 100009415 | Foot Phalanx II; Metatarsal II; Hand Phalanx IV; Metacarpals II, V; |
|  | 100018494 | 100022360 | Foot Phalanges I-III, V; Metatarsals I-III, V; Hand Phalanges I, III, IV, V; Metacarpals II, III |
|  | 100032576 | 100033757 | Metatarsal V |
|  | 100042238 | 100044322 | Foot Phalanges I-V; Metatarsals I-V; Hand Phalanges I-V; Metacarpals I-V |
|  | 100058266 | 100062127 | Foot Phalanges I-V; Metatarsals I-V; Hand Phalanges I-V; Metacarpals I-V |
|  | 100103369 | 100105117 | Foot Phalanges I-V; Hand Phalanges I-V |
|  | 100110533 | 100116846 | Foot Phalanges I-V; Metatarsals I-V; Hand Phalanges I-V; Metacarpals I-V |
|  | 100124861 | 100126574 | Foot Phalanx I; Metatarsal II; Hand Phalanx III; Metacarpals II, III |
|  | 100131195 | 100133136 | Foot Phalanges I, II; Metatarsals I-V; Hand Phalanges I-V; Metacarpals I-V |
|  | 100140773 | 100142360 | Foot Phalanges I, II; Metatarsals I, II, IV, V; Hand Phalanges I, II, V; Metacarpals I-III, V |
|  | 100196165 | 100197462 | Foot Phalanges III, V; Metatarsals I, IV, V; Hand Phalanges I, III, IV, V; Metacarpals I-III, V |
|  | 100199818 | 100203168 | Foot Phalanges I-V; Metatarsals II-V; Hand Phalanges I-V; Metacarpals I-V |
|  | 100224224 | 100226042 | Metatarsals III-V; Hand Phalanx V; Metacarpals II-V |
|  | 100234080 | 100236829 | Metacarpal II, IV, V |
|  | 100241067 | 100243238 | Distal & Proximal Femur; Proximal Humerus; Distal & Proximal Radius; Proximal Tibia; Foot Phalanges I, V; Metatarsals I-V; Hand Phalanx I-V; Metacarpals I-IV |
|  | 100264745 | 100266902 | Foot Phalanges I, II; Metatarsals I, II, IV, V; Metacarpal V |
|  | 100307139 | 100312190 | Distal & Proximal Femur; Distal & Proximal Humerus; Proximal Radius; Proximal Tibia; Foot Phalanges I, II; Metatarsals I, II, IV, V; Hand Phalanx V; Metacarpals II-IV |
|  | 100331806 | 100332722 | Metatarsal V |
|  | 100338262 | 100340520 | Distal Femur; Foot Phalanges I, II, V; Metatarsals I-V; Hand Phalanges I-V; Metacarpals II, III, V |
|  | 100372584 | 100374300 | Distal Femur; Distal Humerus; Foot Phalanges I-III, V; Metatarsals I-V; Hand Phalanges I, II, IV, V; Metacarpals I-V |

The start and end positions of the open chromatin regions were mapped chromosome 15 (*ADAMTS17*) based on human genome assembly GRCh38 (hg38).

We next analyzed *ADAMTS10* and *ADAMTS17* gene expression in human embryonic skeletal elements by comparing normalized read counts as a measure of ADAMTS10 and ADAMTS17 mRNA abundance (Fig. 4D, E). In stylopodial and zeugopodial elements, normalized read counts for *ADAMTS10* were generally higher compared to *ADAMTS17*, suggesting increased expression (Fig. 4D). We did not observe a distinct pattern of *ADAMTS10* or *ADAMTS17* expression based on specific skeletal elements, suggesting that both genes were expressed in autopods of the hind limb and forelimb. Lastly, the overall read counts for *ADAMTS10* and *ADAMTS17* in the stylopod and zeugopod were higher than in the autopods.

Finally, we mapped predicted binding sites of key chondrogenic and osteogenic transcription factors, including SOX9, RUNX2, and ATF4, within 5 kb upstream of the TSS using the Search Motif Tool in the Eukaryotic Promoter Database (Fig. 4F, G)^43^. Overall, transcription factor binding sites upstream of *ADAMTS10* mapped more frequently closer to the TSS (<2 kb) compared to *ADAMTS17*, with the exception of SOX9, SOX6, and ATF4 binding sites. We also noted more SOX9 binding sites upstream of the *ADAMTS17* TSS.

Collectively, these data suggest specific regions of chromatin accessibility in the vicinity of *ADAMTS10* and *ADAMTS17* and correlating with their expression in human embryonic cartilage. In addition, the mapping of chondrogenic and osteogenic transcription factor binding sites in the *ADAMTS10* and *ADAMTS17* promoter region supports the regulation of *Adamts10* and *Adamts17* expression that was observed during chondrocyte hypertrophy.

### *Adamts10* and *Adamts17* inactivation compromised skin development and differentially regulated ECM deposition by skin fibroblasts

*Adamts10*;*Adamts17* DKO skin easily detached and ripped during shaving. We investigated this phenomenon systematically by measuring the thickness of the dermal sub-layers in Masson’s trichrome stained cross-sections (Fig. 5A-E). The overall thickness of *Adamts10* KO and *Adamts17* KO dorsal skin was reduced compared to WT skin and was even further reduced in DKO skin, which was significantly thinner compared to WT and *Adamts10* KO or *Adamts17* KO skin (Fig. 5B). Epidermal layer thickness was slightly but significantly reduced in DKO skin compared to WT and *Adamts10* KO skin but not compared to *Adamts17* KO skin (Fig. 5C, left). The thickness of *Adamts10* KO and *Adamts17* KO dermis and hypodermis was significantly reduced compared to WT and further reduced in the DKO (Fig. 5C, middle, right). The panniculus carnosus (p. carnosus) muscle, which underlies mouse skin, was similarly significantly thinner in *Adamts17* KO and DKO skin sections compared to WT and *Adamts10* KO (Fig. 5D). The proportion of the hypodermis to overall skin thickness was greatly reduced in DKO skin, whereas the proportions of the dermis and p. carnosus were both increased (Fig. 5E). A similar reduction in the hypodermal layer and increase in the dermal layer were evident in *Adamts17* KO skin and to a lesser extent in *Adamts10* KO skin. We also observed a significant reduction in the number of hair follicles in DKO skin compared to all other genotypes, but no significant changes in the *Adamts10* KO or *Adamts17* KO compared to WT skin (Fig. 5F).

**Figure 5.**
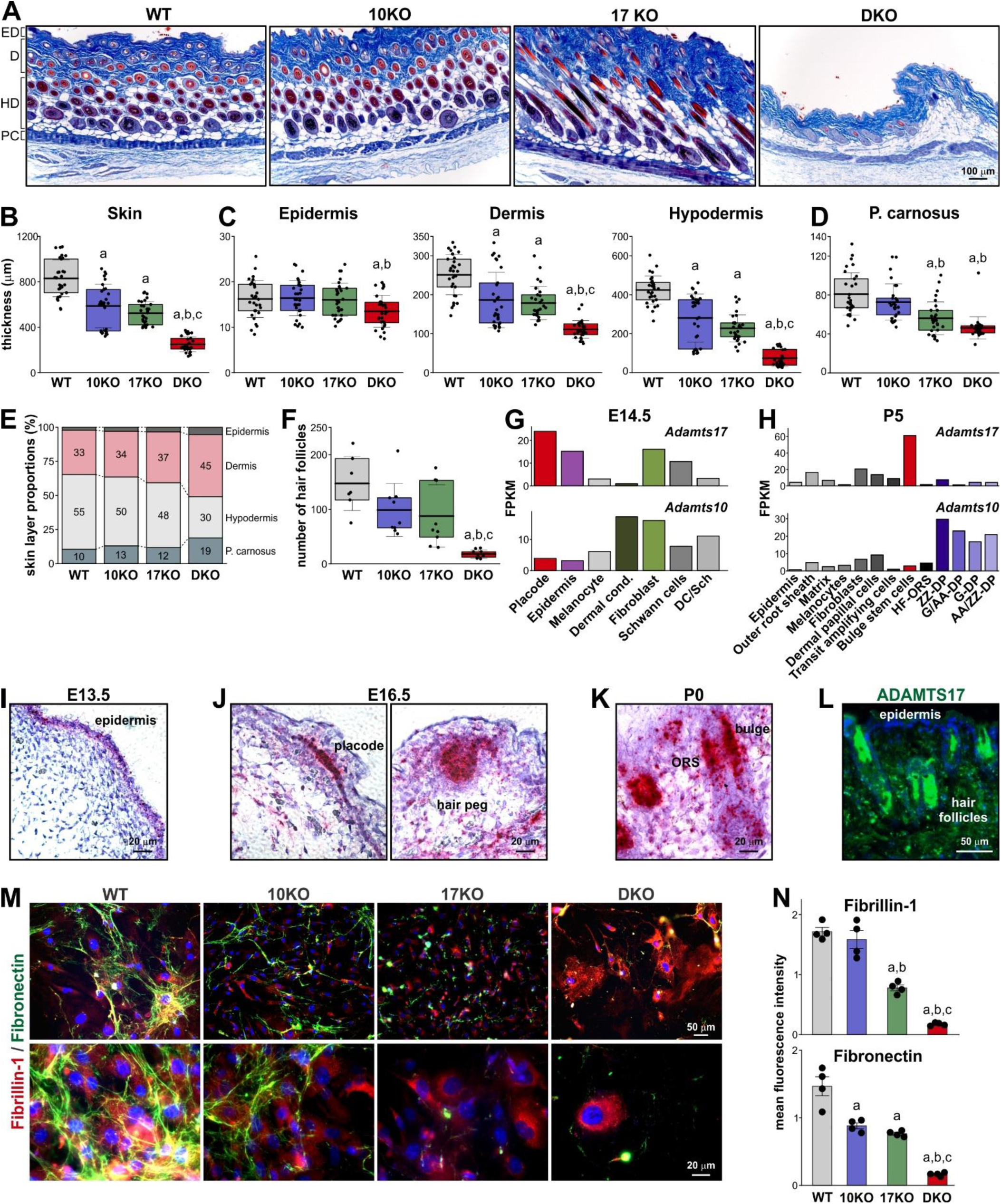
Adamts10;Adamts17 DKO is associated with skin alterations and aberrant ECM deposition by dermal fibroblasts. **A)** Micrographs of Masson’s trichrome-stained cross-sections through dorsal skin from 4-week-old WT, Adamts10 KO (10KO), Adamts17 KO (17KO), and DKO mice. ED, epidermis; D, dermis; HD, hypodermis; PC, panniculus carnosus. **B-D)** Bar graphs showing quantification of overall skin thickness (B) and the thicknesses of the epidermis, dermis, hypodermis (C), and panniculus carnosus (p. carnosus, D). Individual data points represent multiple measurements along the different skin layers from n=3 mice/genotype. **E)** Stacked bar graphs showing the relative proportions of individual skin layers. The percentage values are indicated. **F)** Bar graphs showing the quantification of hair follicle numbers in the skin for each genotype. **G, H)** Bar graphs showing normalized gene expression in fragments per kilobase of transcript per million mapped reads (FPKM) for Adamts10 and Adamts17 in individual skin cell types at E14.5 (G) and P5 (H). Data were extracted from the Hair-GEL database^44,45^. **I-K)** Micrographs showing the localization of Adamts17 mRNA (red/dark purple) in WT skin cross-sections at E13.5 (I), E16.5 (J), and P0 (K) detected by RNAscope in-situ hybridization with a probe specific for Adamts17. Sections were counterstained with hematoxylin. **L)** Micrograph of ADAMTS17 immunostaining (green) of cross-sections through WT skin. Nuclei were stained with DAPI (blue). **M)** Micrographs of primary mouse skin fibroblasts after immunostaining for fibrillin-1 (red) and fibronectin (green). Nuclei were counterstained with DAPI (blue). **N)** Quantification of mean fluorescence intensity from M (n=4 biological replicates). In B, C, D, F, floating bars indicate 25^th^ – 75^th^ percentile range, lines the mean value and whiskers the standard deviation. In N, the bars represent the mean value and the whiskers the standard deviation. Statistical differences in B, C, D, F, N were determined using a one-way ANOVA with post-hoc Tukey test. a, p<0.05 compared to WT; b, p<0.05 compared to Adamts10 KO; p<0.05 compared to Adamts17 KO.

To identify the cellular origins of ADAMTS10 and ADAMTS17 in the skin, we data-mined the Hair-GEL database, which contains single-cell transcriptomic data from E14.5 and P5 skin with a focus on hair follicle development^44,45^. At E14.5, differential expression of *Adamts10* and *Adamts17* was noted in the dermal condensate, where *Adamts10* expression was high, and in the placode and epidermis, where *Adamts17* expression was high (Fig. 5G). In fibroblasts, Schwann cells, or melanocytes, *Adamts10* and *Adamts17* were expressed at similar levels. At P5, *Adamts10* was strongly expressed in subtypes of dermal papilla cells and *Adamts17* was strongly expressed in the bulge stem cell population (Fig. 5H). Expression in other cell types, including fibroblasts, was much lower and differential expression of *Adamts10* and *Adamts17* was less apparent. We next validated temporal *Adamts17* expression dynamics in the skin by RNA in-situ hybridization and immunostaining of embryonic and postnatal mouse skin. At E13.5, *Adamts17* mRNA was localized predominantly in the developing epidermis (Fig. 5I). As previously described, at E16.5, high *Adamts17* mRNA levels were observed in the placode and the hair peg of the developing hair follicles, while lower *Adamts17* expression was observed in the epidermis and cells of the dermal layer (Fig. 5J)^22^. At P0, *Adamts17* mRNA was concentrated in cells surrounding the base of the hair follicle and in the outer root sheath (Fig. 5K). ADAMTS17 immunostaining in postnatal skin confirmed ADAMTS17 protein localization in or around hair follicles and the hair shaft (Fig. 5L). At later postnatal time points up to P21, *Adamts17* mRNA signal was generally low (data not shown).

Finally, we investigated fibrillin-1 and fibronectin ECM deposition in primary skin fibroblasts isolated from WT, *Adamts10* KO, *Adamts17* KO, or *Adamts10;Adamts17* DKO mice. Strikingly, skin fibroblasts from DKOs did not form a fibronectin network and showed abnormal intracellular accumulation of fibrillin-1 (Fig. 5M, N). Fibroblasts isolated from individual *Adamts10* KO or *Adamts17* KO mice showed intermediate phenotypes with a significant reduction of fibronectin in both KOs and a reduction of fibrillin-1 in the *Adamts17* KO. While fibronectin in *Adamts17* KO fibroblasts was present in globular structures or short fibers on the cell surface or in the vicinity of fibroblasts, fibronectin in *Adamts10* KO fibroblasts was largely organized in an extracellular fibrillar network. Notably, in areas of *Adamts10* KO fibroblasts ECM with sparse to no fibronectin network, intracellular fibrillin-1 accumulation was more prevalent.

### Identification of fibronectin and COL6 as ADAMTS17 binding partners

It was previously reported that ADAMTS10 constitutively had poor protease activity due to a degenerated furin cleavage site (GL<u>KR</u> instead of a canonical RX[K/R]R↓ site), which hindered furin- mediated activation (Fig. 6A)^21,31^. In contrast, ADAMTS17 is an active protease based on extensive autoproteolysis at the cell surface^22^. To identify ADAMTS17 substrates, we used an unbiased mass spectrometry (MS)-based N-terminomics strategy, Terminal Amine Isotopic Labeling of Substrates (TAILS). We complemented this approach by yeast-2-hybrid protein-protein interaction screening. For TAILS, we co-cultured human dermal fibroblasts with HEK293 cells stably expressing ADAMTS17 or its active site mutant ADAMTS17-EA (Glu-390 to Ala), which abolishes its autocatalytic activity (Fig. 6B, left)^22^. Following isobaric tag labelling of the samples, we identified differentially abundant N- terminally labeled peptides uniquely present or elevated in ADAMTS17 conditioned medium and prioritized the peptides with neo-N-termini from secreted and/or ECM proteins as candidate substrates (Fig. 6C). Amongst potential ADAMTS17 substrates, we identified peptides from the COL1, COL6A2, and COL6A3 chains, and fibronectin (FN1). Multiple ADAMTS17 peptides were identified in the wild-type ADAMTS17 samples due to its autoctalytic activity and served as positive controls. In parallel, we used the ADAMTS17 ancillary domain (17-AD) as the bait in a yeast-2-hybrid screen to identify binding partners. This approach identified the ECM proteins thrombospondin-1 (THSB1) and fibulin-3 (FBLN3), the secreted proteases ADAM12 and PAPPA, and the extracellular domain of the catalytically inactive receptor tyrosine-protein kinase ERBB3 (Fig. 6D). Most notably, we also identified fibronectin and COL6 as potential ADAMTS17 binding partners, which overlapped with the results from the MS screen. Therefore, we selected fibronectin and COL6 for further investigation and validation as potential ADAMTS17 substrates.

**Figure 6.**
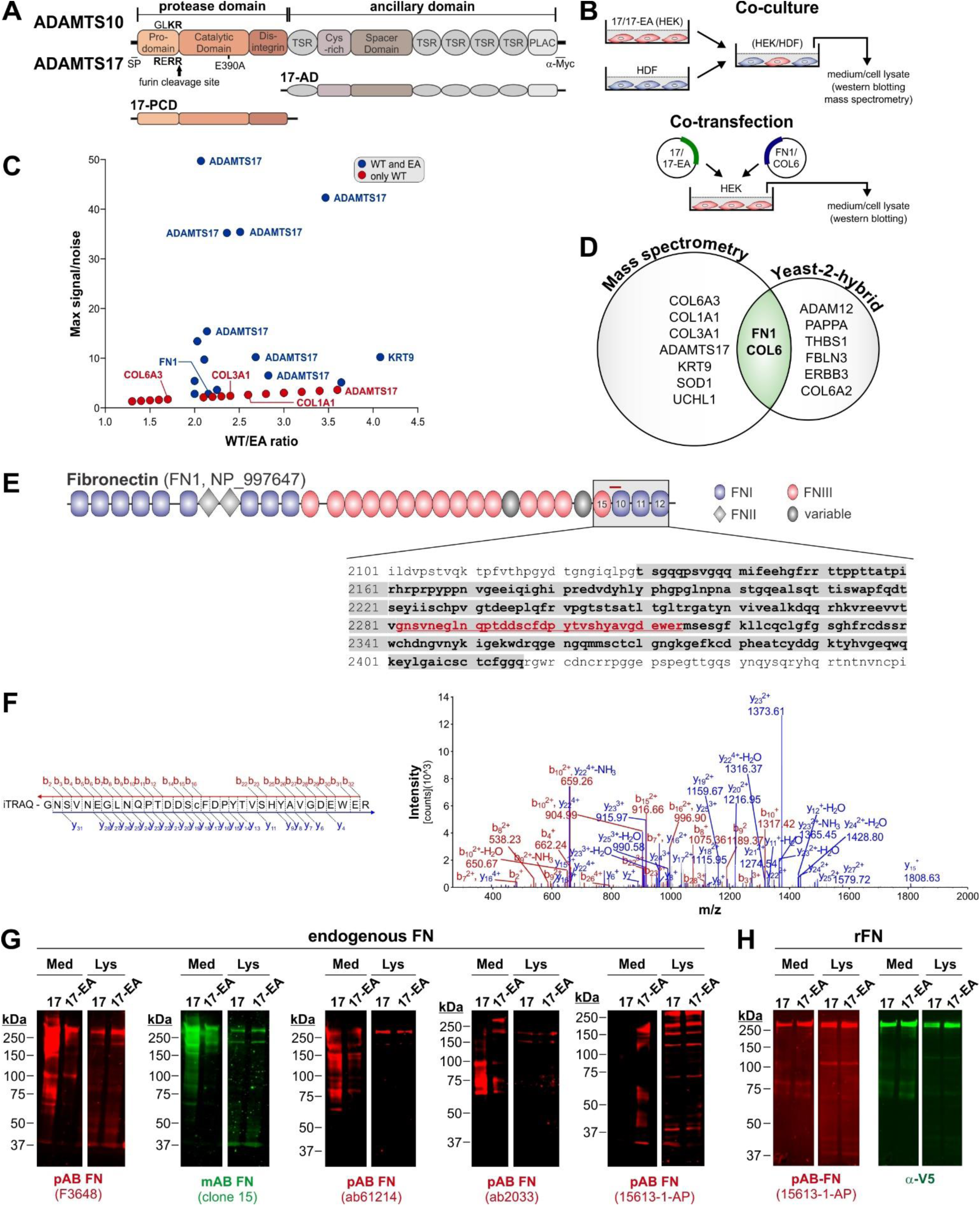
Identification of fibronectin as a potential ADAMTS17 substrate. **A)** Domain organization of ADAMTS10 and ADAMTS17, which are identical. The degenerate (ADAMTS10) and canonical (ADAMTS17) furin processing sites and the localization of the catalytic residue Glu-390 in ADAMTS17 (17) that was mutated into Ala to generate proteolytically inactive ADAMTS17-EA (17-EA) are indicated. The domain organization of the catalytic (17-PCD) and ancillary (17-AD) domain constructs is indicated. **B)** Schematic representation of experimental design for co-culture of human dermal fibroblasts (HDF) with HEK293 cells stably expressing 17- or 17-EA (left) or co-transfection of 17- or 17-EA-encoding plasmids with FN1 or COL6A2-encoding plasmids in HEK293 cells (right). **C)** Volcano plot showing N-terminally labeled peptides identified by N-terminomics method TAILS in conditioned medium from ADAMTS17-expressing HEK293 cells co-cultured with HDFs. Peptides present only in samples from WT ADAMTS17 (red) or enriched in conditioned medium from WT ADAMTS17 co-cultures compared to the co-cultures with proteolytically inactive ADAMTS17-EA suggest ADAMTS17 substrates. **D)** Venn diagram showing overlap of ADAMTS17-cleaved proteins (TAILS) from co-culture systems (left) and binding partners for the ADAMTS17 ancillary domain (17-AD) identified by yeast-2-hybrid screening with a human placenta-derived cDNA library (right). Note that fibronectin (FN1) and COL6 were independently identified in both screens. **E)** Domain organization of fibronectin (FN1, NP_997647) showing the localization of the domains that interacted with 17-AD (grey box, bolded amino acid sequence) and the localization of the peptide identified by TAILS (red bar, red amino acid sequence). **F)** MS2 spectrum of the N-terminally labeled FN1 peptide (GNSVNEGLNQPTDDSCFDPYTVSHYAVGDEWER) showing b and y ions. **G)** Western blot detection of endogenous fibronectin in conditioned medium (Med) and cell lysates (Lys) collected after co-culture of 17 or 17-EA-expressing HEKs with HDFs. A monoclonal (green) and four different polyclonal (red) anti-fibronectin antibodies were used. **F)** Western blot detection of recombinant fibronectin (rFN) in conditioned medium (Med) and cell lysate (Lys) collected after co-expression of 17 or 17-EA with rFN in HEK293 cells. A polyclonal anti-fibronectin antibody (red) and a monoclonal anti V5-tag antibody (green) were used to detect rFN.

Based on the yeast-2-hybrid data, the ADAMTS17 ancillary domain interacted with the C-terminal region of fibronectin comprising FNIII domain #15 and FNI domains #10-12 (amino acid residues 2130 – 2416, NP_997647) (Fig. 6E, F). The N-terminally labeled fibronectin peptide identified in the MS approach localized to the same region (amino acid residues 2282 – 2317) and covered the linker region between FNIII #15 and FNI #10 and the N-terminal part of FNI #10. For biochemical validation of fibronectin as an ADAMTS17 substrate, we first used western blot of conditioned medium and cell lysates collected from human dermal fibroblasts co-cultured with ADAMTS17- or ADAMTS17-EA- expressing HEK293 cells equivalent to the MS approach (Fig. 6B, left). Using five different antibodies against fibronectin, we detected distinct fibronectin fragmentation patterns in conditioned medium in the presence of ADAMTS17-, but not ADAMTS17-EA-expressing HEK293s (Fig. 6G). All antibodies were raised against full length plasma fibronectin, except 15613-1-AP (ProteinTech), which was raised against a region comprising FNIII domain #15 and FNI domains #10-12. Fibronectin fragmentation in the cell lysate, which included the ECM fraction, was not observed. Together, this suggested that ADAMTS17 protease activity correlated with fibronectin fragmentation. To more directly test if ADAMTS17 can cleave fibronectin, we co-transfected HEK293 cells with ADAMTS17 or ADAMTS17-EA- encoding plasmids and a plasmid encoding V5-tagged fibronectin (rFN) (Fig. 6B, right). When we analyzed conditioned medium and cell lysates harvested after co-expression of these plasmids, we did not detect fibronectin fragmentation in the medium or cell lysate with a polyclonal antibody against fibronectin or an antibody against the V5 tag of rFN (Fig. 6H). This suggested that fibronectin may not be a direct ADAMTS17 substrate, or that ADAMTS17 selectively cleaved fibronectin fibrils, which are assembled by dermal fibroblasts.

For COL6A2, the yeast-2-hydrid data suggested an interaction region for 17-AD comprising the C- terminal portion of the central triple helical domain and an N-terminal portion of the von Willebrand factor A (VWA) domain #C1 (amino acid residues 480 – 652, NP_001840.3) (Fig. 7A). TAILS identified an N-terminally labeled peptide originating from COL6A3 (amino acid residues 1348 – 1367, NP_004360.2), which was localized in the C-terminal portion of VWA domain #N4 (Fig. 7B, C). For validation of COL6 as ADAMTS17 substrate, we used the cell culture setups shown in Fig. 6B. In the co- culture system of ADAMTS17-expressing HEKs with HDFs, we did not observe a different pattern of bands originating from endogenous COL6 in conditioned medium in the presence of active ADAMTS17 as detected with a COL6 antibody (Fig. 7D). However, we noticed the disappearance of a ∼125 kDa COL6-reactive band in the cell lysate/ECM fraction when proteolytically active ADAMTS17 was present. When visualizing COL6 ECM deposition in this co-culture system by immunostaining, we observed reduced COL6 staining in the presence of ADAMTS17 compared to ADAMTS17-EA (Fig. 7E, F). We observed a similar difference when fibroblasts were cultured in the presence of cell-free conditioned medium collected from ADAMTS17- or ADAMTS17-EA-expressing HEK293 cells. In the presence of ADAMTS17, COL6 in the ECM was lower than the amount of COL6 deposited in the presence of ADAMTS17-EA (Fig. 7G, H). These observations would be consistent with proteolytically active ADAMTS17 limiting the amount of COL6 deposited in the ECM, potentially via proteolysis, but could also be explained by inactive ADAMTS17-EA promoting COL6 deposition into the ECM. To determine, if COL6 is a direct ADAMTS17 substrate, we co-expressed FLAG-tagged recombinant (r)COL6A2 with ADAMTS17 or ADAMTS17-EA in HEK293 cells. Using western blot detection of the FLAG-tag, we did not observe rCOL6A2 fragmentation in the medium or cell lysate and only detected full length COL6A2 (Fig. 7I). However, we noticed an increase in the band intensity for COL6 in the ADAMTS17-EA lysate, which could be the result of decreased proteolysis or increased cellular retention or cell surface/ECM association. To determine, if ADAMTS17 co-localized with COL6, we cultured fibroblasts producing endogenous COL6, in the presence of the previously described purified recombinant catalytic ADAMTS17 domains (17-PCD) or its ancillary domains (17-AD) and co-immunostained for endogenous COL6 and recombinant ADAMTS17 (α-Myc)^22^. Endogenous COL6 in the ECM of fibroblasts costained with both ADAMTS17 protein constructs, suggesting the possibility of an interaction in the ECM that could be the basis for selective proteolysis of COL6A3 (Fig. 7J). Finally, we analyzed endogenous COL6 distribution and ECM deposition in dermal fibroblasts isolated from a patient with WMS due to a ADAMTS17 Thr343Ala mutation, where we previously showed intracellular retention and reduced ECM deposition of fibronectin, fibrillin-1, and COL1^10^. Compared to control adult human dermal fibroblasts, we observed a strong reduction of COL6 ECM deposition in WMS dermal fibroblasts and concurrent intracellular COL6 accumulation (Fig. 7K, L). This observation was confirmed by western blot analysis, where COL6 was decreased in the medium from WMS patient-derived dermal fibroblasts and increased in the cell lysate compared to adult human dermal fibroblasts (Fig. 7M, N). Collectively, our data suggest that fibronectin and COL6 are potential ADAMTS17 binding partners and/or substrates.

**Figure 7.**
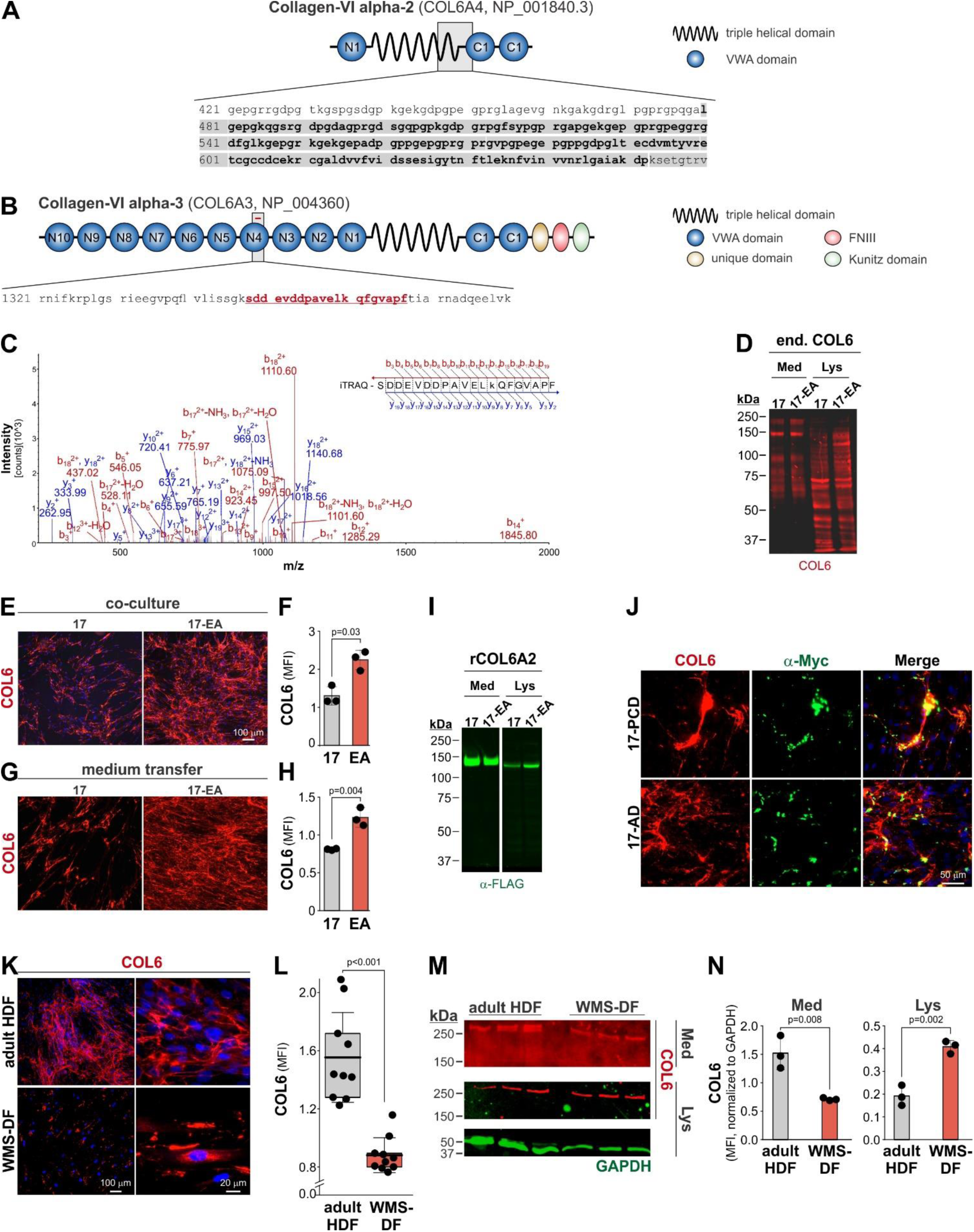
Identification of collagen VI (COL6) as an ADAMTS17 interacting protein and potential substrate. **A)** Domain organization of COL6A2 (NP_0018403) showing the localization of the domains that interacted with 17-AD (grey box, bolded amino acid sequence. **B)** Domain organization of COL6A3 (NP_004360) showing the localization of the peptide identified by MS (red bar, red amino acid sequence). **C)** MS2 spectrum of the N-terminally labeled ADAMTS17-digested COL6A3 peptide (SDDEVDDPAVELkQFGVAPF) showing b- and y-ions. **D)** Western blot of endogenous (end.) COL6 (red) in conditioned medium (Med) and cell lysate (Lys) collected from co-cultures of 17 or 17-EA-expressing HEK293 cells with HDFs. **E)** Micrographs of endogenous COL6 deposition (red) in the ECM of HDFs co-cultured with 17- or 17-EA-expressing HEK293 cells. Nuclei were stained with DAPI (blue). **F)** Quantification of the mean fluorescence intensity of the COL6 signal (n=3 replicates). **G)** Micrographs of endogenous COL6 deposition (red) in the ECM of HDF after culture in the presence of conditioned medium from 17- or 17-EA-expressing HEK293 cells. Nuclei were stained with DAPI (blue). **H)** Quantification of the mean fluorescence intensity of the COL6 signal (n=3 replicates). **I)** Western blot of recombinant COL6 (rCOL6) in conditioned medium (Med) and cell lysate (Lys) collected after co-expression of 17 or 17-EA and rCOL6A2 in HEK293 cells using a monoclonal anti FLAG-tag antibody (green). **J)** Micrographs of HDFs cultured in the presence of 50 μg/ml of purified recombinant 17-PCD and 17-AD protein (see Fig. 5A for domain organization) co-stained for endogenous COL6 (red) and the Myc-tag of the recombinant ADAMTS17 protein fragments (green). Nuclei were stained with DAPI (blue). **K)** Micrographs of adult HDFs and WMS-patient-derived dermal fibroblasts (WMS-DF) for endogenous COL6 (red). Nuclei were counterstained with DAPI (blue). **L)** Quantification of the mean fluorescence intensity of the COL6 signal (n=3 replicates, 2-3 fields-of-view). **M)** Western blot of endogenous COL6 (red) and GAPDH (green) in conditioned medium (Med) and cell lysate (Lys) collected from HDF and WMS-DF cultures. **N)** Quantification of COL6 band mean fluorescence intensities normalized to GAPDH. In E, G, M, bars represent the mean value and whiskers the standard deviation. In K, the floating bars indicate the 25^th^ – 75^th^ percentile range, the lines the mean value and whiskers the standard deviation. Statistical differences in E, G, K, M were determined using a two-sided Student t-test.

## Discussion

The fact that mutations in *ADAMTS10* and *ADAMTS17* can both cause short stature in WMS unequivocally implicates both genes in the regulation of bone growth via an impact on growth plate function. Since it is unclear if and how both genes interact or cooperate in this process, we analyzed the phenotypes of *Adamts10*;*Adamts17* DKO mice. We showed that combined inactivation of ADAMTS10 and ADAMTS17 exacerbated bone shortening when compared to individual KOs with ADAMTS17 gene dosage having an apparently stronger effect and compromised postnatal survival. In addition, we showed that ADAMTS10 and ADAMTS17 are required for skin development, which may relate to stiff and thickened skin described in WMS patients^3,46^. Finally, we identified fibronectin and COL6 as potential ADAMTS17 substrates in high-throughput unbiased screens. This could point towards a potentially distinct function for ADAMTS17 as a regulator of ECM formation and homeostasis. Indeed, such a function is known for ADAMTS10, specifically in the enhancement of fibrillin-1 assembly, since endogenous ADAMTS10 is only slightly activated by furin^21^.

Bone growth is largely driven by chondrocyte proliferation in the growth plate and their subsequent hypertrophic expansion. The disruption of either process can result in bone shortening and short stature^37,47–49^. The reduction in the width of the hypertrophic zone observed in the DKO growth plate could thus be attributed to decreased formation and/or accelerated turnover of hypertrophic chondrocytes, which depend on chondrocyte proliferation and matrix metalloprotease (MMP)- mediated ECM remodeling at the ossification front, respectively^50–52^. Since we did not observe changes in the dimensions of the proliferative zone, we suggest that hypertrophic *Adamts10*;*Adamts17* DKO chondrocytes turnover faster. Hypertrophic chondrocyte turnover requires the transition of a cartilage ECM towards a bone ECM that is primed for mineralization. Key enzymes that regulate these processes include MMP13 and MMP9, which can cleave COL2 and COL10 or promote the vascularization of the growth plate^51,52^. When MMP13 or MMP9 were inactivated, the length of the hypertrophic zone in developing bones was increased by ∼70% and primary ossification was delayed, indicating a delay in chondrocyte turnover, in particular chondrocyte apoptosis^51,52^. Since we observed a shorter hypertrophic zone, we postulate that ADAMTS10 and ADAMTS17 may attenuate chondrocyte turnover potentially by regulating the activity of MMP13 or MMP9^53^. In this context, it is interesting to note that ADAMTS10 WMS knock-in and ADAMTS17 KO mice show opposite growth plate phenotypes, each resulting in reduced bone length. The knock-in of the ADAMTS10 Ser236X mutation (Arg237X in humans) resulted in an expansion of the hypertrophic zone, which correlated with a 6-10% reduction in long bone length^34^. In contrast, inactivation of *Adamts17* resulted in a shorter hypertrophic zone, which also correlated with a 6-10% reduction long bone length^33^. In *Adamts17* KO mice, however, chondrocyte proliferation or apoptosis was not changed^33^. These reports would suggest that ADAMTS10 and ADAMTS17 have distinct roles in regulating growth plate activity and chondrocyte hypertrophy, both resulting in shorter bones. This interpretation is further supported by our findings reported here, that maturation and mineral deposition in ADAMTS17-deficient primary chondrocytes was strongly reduced, but unaffected in ADAMTS10-deficient chondrocytes. The latter findings could also be explained by differential compensation, where ADAMTS17 can compensate for the lack of ADAMTS10 in primary chondrocytes, but not vice versa. As an alternative explanation, ADAMTS10 and ADAMTS17 could regulate signaling pathways that drive growth plate activity and bone growth. In this context, it was shown that reduced BMP signaling in *Adamts17* KO mice delayed terminal differentiation of chondrocytes^33^. In the skin of *Adamts10* WMS knock-in mice, BMP signaling but not TGFβ signaling was similarly reduced^34^. How ADAMTS10- or ADAMTS17-deficiency translates into reduced BMP signaling is not clear, but could be secondary to changes in fibronectin and fibrillin-1 deposition, which regulates extracellular BMP activity^54,55^. Taken together, our data and data from others are consistent with ADAMTS10 and ADAMTS17 regulating different aspects of growth plate activity with ADAMTS17 having a stronger effect.

Skin thickening is a feature of WMS syndrome, and decreased skin elasticity in *Adamts10* KO and *Adamts17* KO mice as well as increased skin thickness in *Adamts10* WMS knock-in mice were reported previously^31,33,34^. In contrast, we noticed fragile skin during routine handling of *Adamts10*;*Adamts17* DKO mice and, accordingly, observed reduced skin thickness in *Adamts10*;*Adamts17* DKO mice, which was predominantly driven by a reduction in the hypodermal layer. This suggested that ADAMTS10 and ADAMTS17 play a role in skin development or postnatal homeostasis, with likely partial compensation in the individual KOs given the much stronger phenotype observed for the *Adamts10*;*Adamts17* DKO skin. One reason for these contrasting observations could be that skin thickening was previously observed in older mice, i.e. at 3 and 8 months of age in the *Adamts17* KO and at 3 months of age in the *Adamts10* WMS knock-in^33,34^. Therefore, it is possible that the skin thickness might increase in our individual KOs with age. We previously demonstrated that WMS patient-derived dermal fibroblasts harboring an *ADAMTS17* mutation were deficient in secretion and deposition of ECM proteins, in particular fibronectin, fibrillin-1 and COL1^10^. When we examined ECM deposition in primary DKO mouse skin fibroblasts, we also observed compromised ECM deposition. In the absence of both ADAMTS10 and ADAMTS17, fibronectin deposition was almost completely abolished and fibrillin-1 appeared to be retained intracellularly. Individual KOs displayed intermediate phenotypes with a more profound phenotype in *Adamts17*KO fibroblasts and a somewhat milder phenotype in *Adamts10*KO fibroblasts. In *Adamts17*KO fibroblasts, fibronectin was deposited as globular speckles, but did not form elongated fiber-like structures or ECM networks. In *Adamts10*KO fibroblasts, fibronectin did form fibers and ECM networks, but they were less dense compared to WT fibroblasts and showed regions void of fibronectin networks. Interestingly, in these regions intracellular fibrillin-1 immuno-reactivity was more apparent. Collectively, these data support a model where ADAMTS17 regulates the secretion and/or assembly of fibronectin by remodeling fibronectin fibrils, which are required for the formation of stable fibrillin-1 networks, whereas ADAMTS10 plays a more “downstream” role where it could enhance fibrillin-1 assembly as suggested previously^21,41,56^.

Since ADAMTS10 and ADAMTS17 are members of the ADAMTS protease family they are presumed to fulfil their biological function through their respective protease activities. However, ADAMTS10 is the only ADAMTS protease with a degenerated furin cleavage site (GLKR↓, lacking a canonical Arg residue at the P4 position) and is incompletely activated by furin-mediated removal of the inhibitory prodomain^21^. It was shown that ADAMTS10 could cleave fibrillin-1 and fibrillin-2 efficiently after restoring a canonical furin recognition site (RRKR↓) by mutagenesis, suggesting that ADAMTS10 has intrinsic protease activity when activated^21,31^. In *Adamts10* KO and *Adamts10* WMS knock-in mice, fibrillin-2 accumulation in the ciliary zonule of the eye was observed and could be explained by the absence of “fibrillin-ase” activity in ADAMTS10-deficient tissue. Indeed, furin activated ADAMTS10 cleaves fibrillin-2^31^. However, fibrillin-2 accumulation could also be explained by ADAMTS10 promoting the assembly of fibrillin-1, which would be reduced in the *Adamts10* KO and result in increased fibrillin-2 exposure to antibodies in the fibrillin microfibril bundles of the ciliary zonule^21,31,34^. Alternatively, ADAMTS10 could be involved in promoting the switch from developmental fibrillin-2-rich microfibrils to postnatal fibrillin-1-rich microfibrils, which similarly would be lacking or be reduced in ADAMTS10-deficient tissues. Therefore, it is plausible that ADAMTS10 may primarily regulate ECM formation both via protease-dependent and protease-independent activities on fibrillin-2 and fibrilin-1 respectively, modulating the formation and isoform composition of fibrillin microfibrils. For ADAMTS17, we showed that it cleaves itself at multiple sites, including in the active site, and is thus proteolytically active^22^. Therefore, we used a complementary MS and yeast-2-hybrid approach in the quest to uncover ADAMTS17 substrates. In both screens, fibronectin and COL6 were identified as potential substrates and binding partners for ADAMTS17.

In summary, this study provides evidence that ADAMTS10 and ADAMTS17 may operate in distinct ways in the pathways that regulate bone growth and skin development through regulation of ECM deposition and turnover and provide additional understanding of the mechanisms underlying WMS.

## Materials and Methods

### Mouse models

*Adamts10* KO mice generated by Deltagen Inc. and their genotyping were described previously^31^. In brief, 41 bp of *Adamts10* exon 5 were replaced with an IRES-lacZ-neo cassette resulting in a frameshift and a premature stop codon triggering nonsense-mediated mRNA decay of the *Adamts10* mRNA. The mice were licensed from Deltagen Inc. (agreement #AGR-17486) and maintained in the C57BL/6 strain. *Adamts17* KO mice were generated in C57BL/6 ES cells by CRISPR/Cas9-induced non-homologous end-joining mutagenesis by contract to Applied StemCell, Inc. A guide RNA targeting *Adamts17* exon 3 (5’-GTCCCTCCACCTCCGTAGCA-3’) was co-injected with Cas9 protein into blastocysts to generate F0 mice, which were screened for frameshift-causing indels. F0 mice harboring an AT insertion in *Adamts17* exon 3 were identified by Sanger sequencing and used to generate F1 mice with the germline-transmitted *Adamts17* AT insertion. F1 mice harboring the AT insertion were identified by Sanger sequencing of a PCR product generated with primers adjacent to exon 3 (F primer: 5’-GAGAGCATCTGATCAGACGCAAATGG-3’, R primer: 5’-CATGTGACCCACAGAGTGTCAGC-3’). *Adamts17* KO mice were rederived at the Icahn School of Medicine. *Adamts17* Het and KO mice were genotyped using DNA from clipped toes as template for a PCR reaction with the following primers: TS17-F 5’-CAGCAGACAGAAGCACAAGCC-3’ and TS17-R 5’-TAGAATCATGGCCCTGACACC-3’. The resulting PCR product was isolated and submitted for Sanger sequencing using the TS17-F primer (Psomagen). The *Adamts17* mutant was maintained in the C57BL/6 strain. Mice heterozygous for the *Adamts10* KO and *Adamts17* KO alleles were crossbred to generate *Adamts10* Het;*Adamts17* Het mice, which were then crossbred to generate *Adamts10*;*Adamts17* DKO mice. *Adamts10* Het or *Adamts17* Het mice were crossbred to generate age- and sex-matched WT, *Adamts10* KO, or *Adamts17* KO mice. Mice were used between 4-8 weeks of age and data from both sexes were combined. All mouse experiments were approved by the Institutional Animal Care and Use Committee (IACUC) of the Icahn School of Medicine at Mount Sinai (protocol numbers IACUC-2018-0009, PROTO202000259, and TR202300000105).

### X-ray imaging

After sacrifice, intact mouse limbs were imaged using a high-resolution radiographic system (UltraFocus digital X-ray cabinet, Faxitron Bioptics). A 10 mm metal rod was used as a scale to enable quantification of bone length.

### Histology

Limbs from 4-week-old mice were disarticulated and the tibia and femur cut mid-shaft to dissect the knee joint. Muscle and other soft tissue were removed and the knee was fixed in Z-fix (Electron Microscopy Science) for 48 h. After decalcification in 14% EDTA solution, the knee joints were dehydrated, embedded in paraffin and sectioned through the middle of the knee. Sections were then stained with hematoxylin & eosin (H&E) for histomorphometry. Proximal tibial growth plates were imaged and the dimensions of the growth plate regions measured at multiple points across the growth plate using ImageJ Fiji (NIH)^57,58^. Dorsal skin was first shaved to remove hair, and a full thickness rectangular strip was dissected and flattened on filter paper. The filter paper with the skin was then immersed in Z-Fix Zinc Formalin Fixative for 24h, processed and paraffin embedded. Cross sections were stained with Masson’s trichrome stain and imaged. Skin thickness and the thickness of individual skin layers was measured from micrographs using ImageJ Fiji.

### Cell culture assays

Human embryonic kidney (HEK) 293 cells (CRL-1573) and adult human dermal fibroblasts (HDF, PCS- 201-012) were purchased from ATCC. Isolation and characterization of the WMS patient-derived dermal fibroblasts (WMS-DF) harboring the *ADAMTS17* c.1027A>G (p.Thr343Ala) mutation was described previously^10^. Cells were cultured in Dulbecco’s Modified Eagle Medium (DMEM) supplemented with 10% fetal bovine serum (FBS), 1% L-glutamine, 100 units/ml penicillin and 100 mg/ml streptomycin (PSG) (complete DMEM) in a 5% CO_2_ atmosphere in a humidified incubator at 37°C. Upon reaching confluence, cells were split in a 1:10 (HEK293) or 1:3 (fibroblasts) ratio. Primary fibroblasts were used up to passage 5-7. HEK293 cells stably expressing ADAMTS17 or ADAMTS17-EA were described previously and maintained in complete DMEM supplemented with geneticin (G418)^22^. For co-culture assays, HDFs and 2 × 10^6^ ADAMTS17 or ADAMTS17-EA (4 × 10^6^ cells total) were combined and seeded on a 10 cm cell culture dish. After reaching confluency the cell layer was rinsed with PBS and cultured in serum free DMEM. After 48 h, conditioned medium was collected and proteins were precipitated with 10% trichloroacetic acid. The cell layer was lysed in RIPA buffer (0.1% NP40, 0.05% DOC, 0.01% SDS in PBS). Equal volumes were mixed with 5x reducing SDS loading buffer, boiled at 100 °C and separated via SDS-PAGE for western blotting. For direct ADAMTS17 proteolysis assays, HEK293 cells were co-transfected with ADAMTS17 or ADAMTS17-EA and plasmids encoding FN1 or COL6A2 using Lipofectamine 3000. The plasmid encoding V5-tagged fibronectin was described recently and kindly provided by Dr. Dieter Reinhardt (McGill University, Montreal, Canada)^59^. The COL6A2-encoding plasmid was purchased from Genscript (OHu18654D). After 24 h, cell layers were rinsed with PBS and cultured in serum-free DMEM for an additional 48 h. Conditioned medium was collected and cleared from cell debris by centrifugation. The cell layer was lysed in RIPA buffer, transferred into an Eppendorf tube and cleared of debris by centrifugation. Equal volumes were combined with 5x reducing SDS loading buffer, boiled at 100 °C and separated via SDS-PAGE for western blotting.

### Western blotting

The proteins in equal volumes of media or cell lysates were separated on polyacrylamide gels using SDS-PAGE and blotted onto polyvinylidene difluoride (PVDF) membranes (Immobilon-FL, Merck Millipore Ltd.) using the semi-dry Bio-Rad trans-Blot® Turbo transfer system for 33 min at 25 V (Bio- Rad) or a wet transfer system for 1.5 h at 70 V at 4 °C both operated with 25 mM Tris, 192 mM glycine, 20% methanol as transfer buffer. Membranes were blocked with 5% (w/v) milk in TBS (10 mM Tris-HCl, pH 7.2, 150mM NaCl) for 1 h at RT and incubated with the following primary antibodies diluted in 5% (w/v) milk in TBS-T (TBS + 0.1% Tween 20) at 4 °C overnight: polyclonal antibodies against fibronectin (F3648, 1:1,000, Sigma; ab2033, 1:200, Millipore; 15613-1-AP, 1:1000, ProteinTech; ab61214, 1:200, Abcam), monoclonal antibody against fibronectin (cl 15, 1:400, Sigma), polyclonal antibody against COL6 (ab6588, 1:500, Abcam), monoclonal antibodies against the V5-tag (mAB, 1:500, Invitrogen) and the FLAG-Tag (M2, 1:500, Sigma), or a monoclonal antibody against GAPDH (1:1,000, Millipore). After incubation with the primary antibody, membranes were rinsed with TBS-T 3 × 5 min at RT and incubated with IRDye-goat-anti-mouse or goat-anti-rabbit secondary antibodies (1:10,000 in 5% (w/v, Jackson ImmunoResearch Laboratories) in TBS-T for 2 h at RT. Membranes were then rinsed 3 × 5 min with TBS-T, once in TBS and imaged using an Azure c600 Western blot imaging system (Azure Biosystems). Band intensities were quantified using the AzureSpot analysis software and normalized to GAPDH.

### Immunostaining of tissue sections and cell layers

Growth plate and skin sections were de-paraffinized and rehydrated followed by protease-mediated antigen retrieval using HistoZyme (Diagnostic BioSystems). After blocking with 5% BSA in PBS, sections were incubated with a mouse monoclonal antibody against ADAMTS17, which was raised against human ADAMTS17 (3B7, 1:500, Novus Biologicals) in blocking buffer in a humidified chamber overnight at 4 °C. Sections were rinsed in PBS 3 × 10 min at RT and incubated in AlexaFluor 488-labeled secondary goat-anti-mouse antibody in blocking buffer at for 1 h at RT. Slides were cover-slipped with ProLong Antifade Gold mounting medium and imaged using a Zeiss Axio Observer Z1 Fluorescence Motorized Microscope w/Definite Focus.

HEK293 cells and HDFs or co-cultures thereof, or primary chondrocytes were seeded in 8-well chamber slides (50,000 cells/well) (Celltreat Scientific Products) and cultured in complete DMEM for 3-4 days. For the analysis of COL6, complete DMEM was supplemented with 100 μM ascorbic acid (ThermoFisher). After medium removal, cell layers were rinsed with PBS and fixed for 5 min with 150 μl ice-cold 70% methanol/30% acetone (Thermo-Fisher), which permeabilized the cells, or with 4% PFA for 20 min to only stain cell surface and ECM proteins, followed by permeabilization with 0.1% Triton X- 100 prior to incubation with the secondary antibody. After fixation, cells were rinsed with PBS and blocked with 10% normal goat serum (Jackson ImmunoResearch Laboratories) in PBS for 1 h at RT. For Okadaic acid (Sigma Aldrich #459620) chondrocytes were treated with 50nM Okadaic acid and the controls were treated with DMSO for 24 hours and allowed to differentiate. For immunostaining cells were fixed with 4% PFA without permeabilization. Cells were incubated with primary antibodies against fibrillin-1, fibronectin, COL6, anti-Myc-tag, or ADAMTS17 diluted in blocking buffer overnight at 4 °C. Cells were rinsed 3 × 5 min with PBS and incubated with AlexaFluor labeled secondary goat-anti- mouse or goat-anti-rabbit antibodies (1:350 in blocking buffer) (Jackson ImmunoResearch Laboratories) for 2 h at RT followed by 3 × 5 min rinses with PBS and mounting with ProLong Gold Antifade Reagent with DAPI (Thermo-Fisher). Slides were imaged using a Zeiss Axio Observer Z1 Fluorescence Motorized Microscope w/Definite Focus and Zeiss Zen Microscope Software and ImageJ Fiji were used to quantify fluorescence pixel intensities.

### 10T1/2 pellet culture for chondrocyte-like differentiation

C3H/10T1/2 cells (CCL-226) were purchased from ATCC and cultured according to ATCC’s instructions in Eagle’s Basal medium^60,61^. For maintenance, cells were sub-cultured prior to confluence. For pellet cultures, C3H/10T1/2 cells were detached with trypsin-EDTA and 5 × 10^5^ cells/ml were centrifuged at 500 g for 5 min in 15-ml tubes and cultured in serum-free DMEM supplemented with 0.1 μM dexamethasone, 0.2 mM L-ascorbic acid-2 phosphate, insulin-transferrin-selenium supplement and 10 ng/ml TGF-β1. The medium was changed every 48 h.

### mRNA isolation and quantitative real-time PCR

Chondrocyte pellets were removed at the experimental time points, immersed in TRIzol reagent and RNA was extracted according to the manufacturer’s instructions. Pellets were lysed by pipetting the TRIzol up and own several times to ensure complete homogenization. The lysate was collected into sterile tubes and incubated at room temperature for 5 min. Following lysis, 0.2 ml chloroform was added per 1 ml of TRIzol, the Eppendorf tubes were inverted several times and incubated at RT for 2-3 minutes. To separate the organic and inorganic phase, samples were centrifuged at 12,000 g for 15 min at 4 °C. The aqueous phase containing RNA was carefully transferred to a new Eppendorf tube and the RNA was precipitated by adding 0.5 ml of isopropanol per 1 ml of TRIzol reagent and samples were incubated for 10 min at RT. The RNA was pelleted by centrifugation at 12,000 g for 10 min at 4 °C. The supernatant was discarded and the RNA pellet was washed with 1 ml 75% ethanol per 1 ml of TRIzol, followed by centrifugation at 7,500 g for 5 min at 4 °C. The RNA pellet was air-dried for 5-10 min at RT and dissolved in 30 – 50 μl of RNase-free water depending on pellet size. RNA concentration and purity were measured using a Nanodrop OneC spectrophotometer (ThermoFisher). RNA preparations were further purified using DNAse I to remove traces of DNA. Reverse transcriptase was used to convert 1 μg of RNA into cDNA.

Quantitative real-time (qRT)-PCR was performed in triplicates in a 384-well plate format. For each reaction, 2 μl cDNA, 0.5 μl of forward and reverse primers (100 μM stock) and SYBR Green PCR Master Mix (Applied Biosystems) were combined in a total reaction volume of 10 μl. The amplification and detection were performed on an ABI PRISM 7900HT Sequence Detection System (Applied Biosystems). All reactions were run under standard cycling conditions. The following PCR primer pairs were used: Gapdh (F: 5’-AGGTCGGTGTGAACGGATTTG-3’, R: 5’GGGGTCGTTGATGGCAACA-3’ or F: 5’- CTTTGTCAAGCTCATTTCCTGG-3’, R: 5’-TCTTGCTCAGTGTCCTTGC-3’), Adamts10 (F: 5’-CCCGCCTATTCTACAAGGTGG-3’, R: 5’- GCCTTCCCGTGTCCAGTATTC-3’ or F: 5’-GTAGTGGAGTGCCGAAATCAG-3’, R: 5’-CAGCGTGACCAGTTTCCTAC-3’), Adamts17 (F: 5’-CTGCTGTATTTGTGACCAGGAC-3’, R: 5’- AGCACACATTTCCTCTTAGCAC-3’ or F: 5’-CCTTTACCATCGCACATGAAC-3’, R: 5’-ATTCCGTCCTTTTACCCACTC-3’), Fbn1 (F: 5’-GGACAGCTCAGCGGGATTG-3’, R: 5’- AGGACACATCTCACAGGGGT-3’), *Fn1* (F: 5’- GCTCAGCAAATCGTGCAGC -3’, R: 5’- CTAGGTAGGTCCGTTCCCACT -3’), *Col2a1* (F: 5’- GGGAATGTCCTCTGCGATGAC-3’, R: 5’-GAAGGGGATCTCGGGGTT-3’), *Col10a1* (F: 5’- TTCTGCTGCTAATGTTCTTGACC-3’, R: 5’- GGGATGAAGTATTGTGTCTTGGG-3’).

### Isolation and differentiation of primary chondrocytes

Primary costal chondrocytes were isolated as described previously^62^. The ribcage of P5 mouse pups was dissected, flattened and soft tissue was removed. The cartilaginous portions of the ribs were then transferred into PBS containing 10× penicillin and streptomycin (Gibco). The ribs were digested with 15 ml pronase (2mg/ml, Sigma-Aldrich) in a 50 ml Falcon tube for 1 h at 37 °C in a tissue culture incubator under a 5% CO_2_ atmosphere. The pronase solution was replaced with 15 ml of collagenase D (3 mg/ml, Sigma-Aldrich) followed by incubation for 1 h under the same conditions with gentle agitation every 10 min. After 45 min of incubation, the collagenase D solution was vigorously pipetted up and down over the thoracic cages. The solution was finally transferred into a 50 ml falcon tube. The soft tissue debris was removed as it sediments slower and the process was repeated with sterile PBS buffer. The cleaned cartilage was then digested again with 15 ml collagenase D solution for 4-5 hours at 37 °C in the cell culture incubator. Finally, primary chondrocytes were released by pipetting the solution containing the cartilage fragments up and down ∼10 times. This digest was passed through 40 μm cell strainer and pelleted by centrifugation at 300 g for 5 min. The chondrocyte pellet was rinsed twice with PBS and chondrocytes were resuspended in 10 ml complete DMEM medium, plated at a density of 10^5^/cm^2^ on 6-well plates and cultured in complete DMEM. Upon reaching confluency, the medium was replaced with chondrocyte maturation medium (complete DMEM supplemented with 50 μg/ml ascorbic acid and 10 mM β-glycerophosphate). The medium was changed every 2 days and mineral deposition was visualized after 21 d by alizarin red staining. For this, the differentiated and matured chondrocytes were rinsed with PBS and fixed with 10% formaldehyde (MP Biomedicals) for 15 minutes at RT. The cell layer was rinsed twice with distilled water and incubated in 1 mL of 40 mM alizarin red staining solution (pH 4.1) per well for 20 min at RT under gentle agitation on a shaker. Plates were incubated at room temperature for 20 minutes with gentle shaking. The alizarin red staining solution was removed and the wells were rinsed several times with 5 mL of distilled water. Prior to brightfield imaging, excess water was removed and the cell layer was dried at RT. The cell layers were imaged and the stained mineral deposition was quantified using the ImageJ Fiji.

### ATAC-seq and transcriptomics analyses of ADAMTS10 and ADAMTS17 expression

Sample generation, ATAC-seq, RNA sequencing and data analyses for the data set have been described recently^42^. No additional human products of conceptions that were not previously described, were used for this manuscript. In brief, epiphyses (long bones) or whole elements (pooled phalanges or metapodials) were micro-dissected and RNA was extracted after tissue homogenization followed by RNA sequencing. For ATAC-seq, tissues were digested with collagenase to generate single cell suspensions, which were then subjected to the ATAC-seq protocol. The data sets for the stylopodial and zeugopodial elements are deposited in the Gene Expression Omnibus repository under the accession numbers GSE252289 (human long bone skeleton ATAC-seq) and GSE252288 (human long bone skeleton RNA-seq). The data sets for the autopodial elements will be published elsewhere (Okamoto & Capellini, manuscript in preparation).

### RNAScope in-situ hybridization

WT mouse embryos at E13.5, E16.5, and P0 were fixed in 4% paraformaldehyde in PBS overnight at 4°C, dehydrated, and embedded in paraffin. Fresh 6 μm sections were used for in-situ hybridization using RNAscope (Advanced Cell Diagnostics) following the manufacturer’s protocol. Tissue localization of *Adamts17* mRNA was achieved with a probe recognizing the mRNA from mouse *Adamts17* (#316441) in combination with the RNAscope 2.0 HD detection kit “RED”. Tissue sections were counterstained with hematoxylin and cover-slipped with Cytoseal 60 (Electron Microscopy Science). After curation of the mounting medium, sections were observed using an Olympus BX51 upright microscope equipped with a CCD camera (Leica Microsystems) for imaging.

### Isolation of primary mouse skin fibroblasts

Mice were euthanized using CO2 inhalation followed by cervical dislocation and skin fibroblasts were isolated using enzymatic digestion. Following euthanasia, the dorsal skin was shaved and 1-2cm2 section was excised using sterile scissors and forceps. The excised skin was washed in phosphate- buffered saline (PBS). The skin was minced into 1-2mm2 fragments and transferred into a 50ml conical tube. The tissue was digested using 2mg/ml Collagenase type II (Worthington, LS004202) in DMEM. The tube was incubated at 37oC for 1 hour with gentle agitation to release the fibroblasts. After digestion, the cell suspension was triturated with a 10ml pipette to further dissociate the tissue. The suspension was then filtered through a 70um cell strainer to remove undigested fragments. The filtered cell suspension was centrifuged at 300g for 5 minutes at room temperature. The resulting cell pellet was resuspended in DMEM. The cell suspension was plated onto sterile 10cm culture dishes and incubated at 37oC, 5% CO2. 24h post-plating, the medium was replaced to remove non-adherent cells.

### N-terminomics via TAILS

For iTRAQ labelling, we co-cultured HDFs with HEK293 cells expressing ADAMTS17 or ADAMTS17-EA (1 × 10^6^ cells each, two 10 cm dishes). After reaching confluence, the cell layer was rinsed with PBS and phenol-free serum-free DMEM was added. After 2 d and 4 d, the medium was collected and EDTA (10 mM final concentration) and one tablet of Complete EDTA-free protease inhibitors (Roche), dissolved in 250 μl water, were added. The medium was cleared of cellular debris by centrifugation at 500 rpm for 5 min, sterile filtrated through a 0.22 μm filter, and stored at -80 °C. Media (∼40 ml) were thawed and concentrated to 2.5 mL with centrifugal filter devices (Amicon Ultra-4, 3kDa cut-off; #UFC800324). Proteins were then precipitated by adding 20 mL of ice-cold acetone and 2.5 mL ice cold methanol followed by vortexing and incubation at -80 °C for 3 h. Protein precipitates were collected by centrifugation at 14,200 g in a Beckman JS-13.1 outswing rotor at 4 °C for 20 min. The supernatant was decanted and the protein pellets were air dried. Air-dried pellets were dissolved in 360 μL 8 M guanidine-HCl, 504 μL double-distilled water, and 288 μL 1M HEPES buffer resulting in terminal amine isotopic labeling of substrates (TAILS) buffer (final concentration: 2.5 M GuHCl, 250 mM HEPES, pH7.8).

300 µg of protein from the TS17X1-WT and TS17X1-mut samples were prepared for iTRAQ-TAILS as previously described^63^. In brief, protein was reduced and alkylated with DTT and IAA respectively. Proteins samples were mixed at a 1:1 ratio with iTRAQ labels 113 (TS17X1-WT) or 115 (TS17X1-mut) in DMSO at 37° C overnight and quenched with 1 M tris pH 8. Samples were then combined prior to overnight digestion with trypsin. N-terminally labeled peptides were enriched as previously described using the hyper-dendritic polyglycerol aldehyde^64^. Prior to MS analysis, the samples were desalted using a C18 Ziptip and reconstituted in 50 μL 1% acetic acid. Peptides were identified with a Dionex Ultimate 3000 UHPLC interfaced with a Thermo LTQObitrap Elite hybrid mass spectrometer system. The HPLC column was a Dionex 15 cm x 75 μm id Acclaim Pepmap C18, 2μm, 100 Å reversed- phase capillary chromatography column. 5 μL volumes of the extract were injected and the peptides eluted from the column by an acetonitrile/0.1% formic acid gradient at a flow rate of 0.3 μL/min were introduced into the source of the mass spectrometer on-line operated at 1.9 kV. The digest was analyzed using a data dependent MS method acquiring full scan mass spectra to determine peptide molecular weights and product ion spectra to determine amino acid sequence in successive instrument scans. Both collision-induced dissociation (CID) and higher-energy collisional dissociation (HCD) fragmentation methods were performed on the top-8 most abundant precursor ions in each scan cycle. HCD fragmentation is required to quantify the reporter ions of the iTRAQ labels on peptides. The samples were analyzed using a data dependent acquisition (DDA) using both HCD and CID fragmentations. The resulting data were searched using Sequest program which integrated in Proteomics Discoverer 2.5 software package against Uni-prot human protein sequence database (March, 2024). Trypsin (semi) was set as protease, carbomidomethylation of Cys was set as a static modification and iTRAQ 8-plex of peptide N-terminal, Lys and Tyr, oxidation of Met, and N-term Gln cyclization were set as dynamic modifications.

### Yeast-2-hybrid screening

An ULTImate Y2H SCREEN was performed by Hybrigenics Services using ADAMTS17-AD (aa 546-1122) as bait together with the Prey Library Human Placenta_RP6. The bait was cloned into the pB27 (N-LexA-bait-C fusion) vector. 60 clones were processed and 170 million interactions analyzed.

### Statistical analyses

Statistical analyses were performed using the OriginPro 2018 software. N=2 samples were compared with a two-sided Student’s t-test and n≥3 samples with a one-sided ANOVA. P-values <0.05 were considered statistically different.

## Author contributions

NT, SZK, ZB, LWW, BBW and DRM, performed experiments and analyzed data. SSA and DH analyzed data and contributed to study design. DR, ASO, TDC analyzed ATAC-seq and transcriptomics data. DH conceptualized the study, performed experiments, analyzed data, drafted the manuscript and prepared the figures. All authors edited the manuscript and figures and approved the final version.

## Acknowledgements

This research was in part supported by a grant from the National Institutes of Health (R01AR070748 to D.H.). We thank Dr. Dieter Reinhardt (McGill University, Montreal, Canada) for kindly providing the fibrillin-1 antibody and the V5-tagged rFN1 expression plasmid. We thank Damien Laudier (Orthopedic Research Laboratories, Icahn School of Medicine at Mount Sinai, New York, NY) for growth plate and skin histology.

## Data Availability

The data sets for the stylopodial and zeugopodial elements are deposited in the Gene Expression Omnibus repository under the accession numbers GSE252289 (human long bone skeleton ATAC-seq) and GSE252288 (human long bone skeleton RNA-seq). The data sets for the autopodial elements will be published elsewhere (Okamoto & Capellini, manuscript in preparation) and made publicly available.

